# Mercury bioaccumulation varies across breeding-site height and life-history traits in Amazonian phytotelm-breeding frogs

**DOI:** 10.64898/2026.10.13.758764

**Authors:** Bibiana Rojas, Paul Kesseler, Lia Schlippe Justicia, Andrius Pašukonis, Joan Espígol, Doriane Hagnier, Shubham Soni, Valentine Alt, Jérémy Lemaire, Valeria Marasco

**Author notes:** Shared last authorship (Jérémy Lemaire, Valeria Marasco).

## Abstract

Mercury (Hg) concentrations in the environment have significantly increased in the Amazon. Despite the observed deleterious effects on freshwater wildlife, the impacts of Hg contamination in phytotelmata (i.e., waterbodies contained in plant structures) and the species therein remain undocumented. We investigated the effects of Hg contamination on a phytotelm-breeding frog community in an otherwise pristine rainforest in French Guiana. Total Hg concentrations were measured in 28 pools at different heights (i.e. ground to canopy), and in the bodies of 103 tadpoles from five species with contrasting life histories. Tadpoles from ground pools had higher Hg concentrations than those from arboreal pools, revealing a vertical gradient in Hg contamination; tadpoles from species with slow development and parental care (i.e., tadpole transport) exhibited higher Hg concentrations than those of species with fast development without parental care. Finally, although Hg burdens were not associated with oxidative DNA levels, individuals reaching metamorphosis exhibited lower DNA damage than early-stage tadpoles, possibly reflecting reduced oxidative damage resulting from cellular remodelling undergone during metamorphosis, or selective mortality processes. Our findings highlight the joint influence of habitat type and life-history traits on Hg accumulation and that, even in pristine forests, Hg exposure varies strongly across microhabitats and species.

## Introduction

Artisanal and Small-Scale Gold Mining (ASGM) is the largest anthropogenic global source of both atmospheric and freshwater mercury (Hg)^1^ ASGM is often illegal and tends to occur in remote places, frequently in or next to protected areas and biodiversity hotspots^2^, making the impacts of Hg contamination on local wildlife difficult to assess. Intact forest canopies capture substantial quantities of atmospheric Hg^3^, ^4^, which initially accumulates on foliage and other canopy surfaces, and is later transferred to the forest floor via throughfall and litterfall4-6. Consequently, elevated Hg loads accumulate in the upper soil layers in tropical forests^6^, ^7^. Mercury contamination on large bodies of water (i.e., lakes, rivers, streams) has been studied for decades, yet Hg accumulation within forest microhabitats such as phytotelmata remains poorly documented. These small water reservoirs held in plant structures such as bromeliad axils or tree holes^8^, which can occur from the ground all the way to the forest canopy^9^, act as dynamic, natural microcosms that support diverse insect communities and provide essential shelter to numerous species ^10^. Additionally, intrinsic differences in their durability, structure and physical-chemical properties can influence how much Hg is accumulated ^11^ and in what chemical form.

Within ecosystems, inorganic Hg can be transformed into methylmercury (MeHg) through microbial methylation^12^, ^13^. Because of its high bioavailability and tendency to bioaccumulate ^14^, ^15^, MeHg represents the most toxic form of mercury ^12^. Mercury bioaccumulation occurs via the diet, environmental exposure and/or maternal transfer, with negative effects on the organisms’ phenotype, physiology, reproductive success and behaviour ^16^. One of the main mechanisms underlying Hg toxicity is the induction of oxidative stress through increased endogenous production of reactive oxygen and nitrogen species (RONS)^17^. In the absence of repair mechanisms (antioxidants), RONS can inflict substantial damage on macromolecules such as DNA and proteins^18^. The effects of contaminant exposure on organismal health could be especially pronounced and long-lasting if experienced during early development, as physiological alterations in early life can have lasting consequences for individual performance and fitness ^19^.

Most studies on Hg impacts on wildlife have been conducted in temperate ecosystems (but see e.g. ^20^ for exceptions), thus overlooking the environmental characteristics of the tropical zones, the unique features of their wildlife, and the abundance not only of biodiversity hotspots and major wilderness areas, but also of ASGM activities therein. For example, the Amazon basin, a high-biodiversity wilderness area containing almost 10% of the planet’s biodiversity ^21^, ^22^, is a global hotspot for ASGM accounting for approximately 80% of South America’s and 27% of global anthropogenic Hg emissions ^23^. Amazon rainforests also hold one of the world’s highest richness in amphibians 22, with a remarkable diversity of life histories, and reproductive modes ^24^. Among these are semi-terrestrial species that use phytotelmata as both breeding sites for adults and rearing habitats for tadpoles ^25^. Because these habitats are confined, tadpoles are particularly vulnerable to water contamination, making them sensitive bioindicators of ecosystem health ^26^.

With 41% of species having declined or become threatened in the last 60 years, amphibians are currently considered the most endangered vertebrate taxon on the planet ^27^. Threats for Amazonian amphibians are rapidly increasing due to, among other factors, deforestation and the increased exposure to chemical pollutants such as pesticides and heavy metals like Hg ^28^. Recent research has revealed that phytotelmata can have moderate to high Hg levels in the water and dissolved organic matter ^11^. Furthermore, tadpoles can be found in pools with Hg concentrations as high as 8ppm 11, far exceeding the threshold at which freshwater sediments are considered heavily polluted and prone to causing adverse effects in sediment-dwelling organisms (SEL = 2ppm; ^29^). This is problematic considering the potential role of phytotelm-dependent amphibians as pollution vectors of Hg to terrestrial systems upon metamorphosis, and their vulnerability to changes in the environment due to their permeable skin ^30^. Mercury contamination in amphibians may have sublethal effects in adults and larvae, including changes in behaviour^31^, physiology ^32^ and ontogeny ^33^. Yet, despite the predicted negative effects, little is known about how tropical amphibian species respond to Hg exposure in their natural habitats, about the magnitude of Hg contamination and its consequences across species of phytotelm-breeders, and whether such effects differ in relation to key ecological traits.

Here, we investigated the magnitude of mercury contamination and its consequences in a community of phytotelm-breeding amphibians in an otherwise pristine tropical forest area of French Guiana. We examined Hg levels in the water and sediment of pools used for tadpole development, and in tadpoles of five different species with contrasting reproductive ecology and life histories, and which exploit pools at different heights in the vertical gradient of the forest, i.e., from the ground to the canopy. Then, we assessed the levels of oxidative DNA damage present in each species in relation to the amount of Hg accumulated in their body. We expected a positive relation between levels of water Hg and levels of Hg burdens in the tadpoles. Likewise, we predicted higher Hg burdens in tadpoles of (1) species using pools on the ground in comparison to species using pools in the canopy; (2) species with longer larval stages; and (3) more developed individuals, such as those reaching the end of metamorphosis. Consequently, we also predicted higher levels of oxidative DNA damage in individuals (4) with higher Hg burdens and (5) at the more advanced developmental stages.

## 2. Materials and Methods

### 2.1 Study site and species

This study was carried out at Camp Pararé, Nouragues Ecological Research Station, French Guiana (4 02′N, 52 41′W), a site with primary *terra firme* forest about 16Km from known gold mining activities 11. Sampling was done over two field seasons, in April 2024 and between March and May 2025, which fall within the rainy season and within the reproductive season of the studied species (see below).

The studied species were chosen to represent the diversity and complexity of natural history and reproductive ecology of phytotelm-breeders at Nouragues ^34^, including differences in habitat (arboreal or terrestrial) and life history, which ranges from large clutches with no parental care and rapid larval development, to clutch attendance and transport of tadpoles to bodies of water where they complete their (rather slow) development. For this study we focused on the following species, based on their abundance: *Dendrobates tinctorius, Rhinella castaneotica, Osteocephalus oophagus*, *Trachycephalus resinifictrix* and ground-dwelling small dendrobatid species, which include predominantly *Allobates femoralis*, but also *A. granti* and *Ameerega hahneli*. These three species have similar ecologies, use similar aquatic resources and their early stage tadpoles are difficult to discriminate in the field 9. However, because *A. femoralis* is the predominant species in our study area, and hence the most likely representative of these species, we are hereafter referring only to *A. femoralis*.

*Dendrobates tinctorius* and *A. femoralis* are terrestrial, diurnal poison frog species from the families Dendrobatidae and Aromobatidae, respectively ^35^. In both species, males perform parental care, which consists of clutch attendance and tadpole transport to phytotelmata, where they stay unattended until metamorphosis. *D. tinctorius* is commonly found near canopy gaps at elevations ranging from 0 to 600 m.a.s.l. ^36^ and uses a wide variety of phytotelmata, from the forest floor to the canopy (up to 20+ m) for depositing their tadpoles ^9^, ^37^. *D. tinctorius* tadpoles are omnivorous and opportunistic cannibals ^37^, ^38^, and their development until metamorphosis lasts approximately two months ^38^. *D. tinctorius* was declared a protected species in French Guiana in 2020 (Decree No. TREL2032100A, November 19th, 2020). *Allobates femoralis* males use a variety of small terrestrial pools including phytotelmata that are easily accessible from the ground ^9^, ^39^. Tadpoles are omnivorous but feed mostly on detritus ^40^ and their larval development typically lasts between 40 and 50 days ^41^.

*Osteocephalus oophagus* and *Trachycephalus resinifictrix* are two arboreal frogs in the family Hylidae ^42^, which exhibit obligatory and facultative oophagy, respectively. Both species breed in high arboreal phytotelmata, although they can be sporadically found calling and breeding in relatively low arboreal pools ^9^. In *O. oophagus*, clutches range between 10 and 69 eggs, and tadpoles are fully dependent on eggs regularly deposited by the mother at the tadpole rearing site and required for survival beyond early development ^43^. In contrast, *T. resinifictrix* has clutches of up to 1540 eggs, and its tadpoles engage in opportunistic oophagy, consuming conspecific eggs when available but primarily feeding on detritus and other organic material within phytotelmata ^44^. The larval stage lasts approximately 27–50 days in *O. oophagus* ^43^ and 45–75 days in *T. resinifictrix* ^45^.

*Rhinella castaneotica* is a terrestrial toad from the family Bufonidae ^46^. It breeds in forest pools, where females deposit up to 720 eggs which are then left unattended (i.e., no parental care) ^47^. Although there are no reports on the larval diet of *R. castaneotica*, the tadpoles of closely related species are known to be omnivorous grazers, feeding predominantly on algae ^48^. Tadpoles of this species have a remarkably rapid development, with metamorphosis occurring in approximately 20 days (PK, AP, BR pers. obs.).

### 2.2 Data collection in the field

We surveyed predefined working areas by actively searching for phytotelmata containing tadpoles of at least one of the five study species. Rather than following fixed transects, searches were conducted by scanning throughout these study areas, targeting suitable microhabitats such as fallen trees and trees with buttresses or hole structures, typically between 8:30 and 17:30. For each phytotelma (pool, hereon) we collected a 4 ml mixture of water and sediment using a sterile pipette. In addition, we measured pool depth, width, length, and height from the ground to the pool edge, and classified them into six categories: rock pool (water-filled depressions in rocky substrates), palm bract (water accumulated in fallen palm bracts), buttress tree (water retained within buttresses of living trees), fallen log (water-filled cavities in fallen tree trunks), treehole (water-filled holes in trunks of living trees accessible without climbing equipment), canopy pool (water bodies formed in tree structures within the canopy, at heights over 6m and up to 25m). Water pH and temperature were also measured using a pH metre (pH 20, Apera Instruments®). Upon return to the camp at the end of each sampling day, the mixed water–sediment samples were separated into two distinct samples, a sediment sample and a water sample, and stored at - 20°C. This was done to establish which assessment of environmental contamination levels was best related (if any) to the Hg burdens found in tadpoles.

For each pool, we sampled a maximum of five tadpoles per species and pool for the subsequent quantification of Hg concentrations in their tissues. Tadpoles were collected using nets, plastic spoons or Pasteur pipettes whose narrow end had been removed, and photographed on graph paper to subsequently measure its snout-vent length (body length; hereon SVL) and maximum body width using the software ImageJ ^49^. Tadpoles were also weighed to the nearest 0.01 g using a calibrated digital scale, and their developmental stage was determined according to the Gosner scale ^50^ to account for differences in potential Hg exposure duration. Because exact developmental stages could not always be reliably distinguished, we grouped individuals into three developmental categories following Schlippe Justicia et al. (2024): “early” (early pre-metamorphosis, stages 25–29), “late” (late pre-metamorphosis, stages 30–35), and “metamorph” (pro-metamorphosis/metamorphic climax: stages 35-45). When more than five tadpoles of the same species were present, we selected the individuals that maximised the representation of developmental stages across the dataset. Because late-stage tadpoles were encountered less frequently, some pools were revisited after additional developmental time to increase the representation of later-stage tadpoles.

Following collection, tadpoles (body mass < 0.8 g in all cases) were euthanised and subsequently stored in 95% ethanol until laboratory analyses. All individuals were handled with gloves, and all sampling equipment was disinfected with ethanol between pools to avoid cross-contamination and potential pathogen spread. Sample collection was done under the authorisation of the Direction Régionale des Territoires et de la Mer de Guyane (Arrête No. R03-2024-03-26-00001), after evaluation by the CSRPN (Regional Scientific Council for Natural Heritage) and approval by the Scientific Committee of the Nouragues Ecological Research Station.

### 2.3 Mercury quantification

All tadpoles were longitudinally cut into two sections; one half was used for Hg quantification, and the second half for oxidative DNA damage quantification (see below). Ethanol from tadpole samples was evaporated under a nitrogen stream at room temperature. Tadpole samples were cut into small pieces using scissors, then freeze-dried at −80°C for 48h, ground and homogenised. Sediment samples were dried in an oven for 96h at 45°C until they reached a stable weight, then ground and homogenised. Before analysis, the water and sedimentary organic matter samples were defrosted and vortexed for 3 seconds. Total Hg was quantified in all samples by direct measurement using a Direct Mercury Analyzer (DMA-80 evo III, atomic absorption spectrometer, Milestone) at the University of Vienna. Analyses were made on at least two replicates of 0.4 – 7.0 mg dry weight (dw) of each tadpole sample, 1.5-19 mg dw of each sediment sample, and 400 µL of each water mixture sample. Inter-replicate reproducibility for each sample was validated against a relative standard deviation threshold of <10%. The method was validated by analysis of certified reference material (CRM) of TORT-3 (Lobster hepatopancreas, National Research Council of Canada; certified reference Hg concentration: 0.292 ± 0.022 µg.g-1 dw). CRM was analysed at the beginning and end of the analytical cycle, and after every 10 samples. Measured TORT-3 values were 0.292 ± 0.007 µg.g-1 dw (n = 23), with a recovery rate of 100.1 ± 2.3 %. THg concentrations are further expressed in µg.g-1 dw. The detection limit of the DMA-80 was 0.01 ng. Mercury concentrations were below detectable levels in the tissue of six individuals of *R. castaneotica* and one of *A. femoralis*, as well as in the water and sediment of five pools.

### 2.4 Quantification of oxidative DNA damage

We extracted DNA from the individually collected tissue homogenates using DNeasy Blood and Tissue Kit (Qiagen) following the manufactureŕs protocol. The purity and quantity of the DNA extracts was assessed using a NanoDropTM spectrophotometer (NanoPhotometer N60, Implen GmbH). Then, 300 ng of isolated DNA was used to quantify levels of 8-OHdG (8-Hydroxy-2′-deoxyguanosine), which is major product of DNA damage and marker of oxidative stress (Valavanidis et al. 2009), using a competitive immunoassay following the manufacturer recommendations (EpiQuick 8-OHdG DNA Damage Quantification Direct Kit Colorimetric, Epigentek, USA). As the levels of DNA damage in our samples were relatively low, we used a more dilute standard curve (2.5-50 pg µl−1 8-OHdG) than that suggested in the manufacturer’s protocol (5-200 pg µl−1 8-OHdG). DNA damage is expressed as pg of 8-OHdG/μg of DNA. We quantified 8-OHdG in 140 collected tadpoles (out of 164). The intra-assay coefficient of variation averaged 7.4% and the inter-assay coefficient of variation was 17.5%. 8-Hydroxy-2′-deoxyguanosine was below detectable levels in six individuals of *R. castaneotica,* six of *A. femoralis,* seven of *D. tinctorius,* and five of *O. oophagus*.

### 2.5 Statistical analyses

#### Influence of pool characteristics on waterHg and sedimentHg

To examine how pool characteristics influenced the Hg amounts in pool water and sediment we first calculated a proxy for pool’s capacity (volume) using the formula for the volume of a semi-ellipsoid ((4/3*π x pool lengt*ℎ *x pool widt*ℎ *x pool dept*ℎ)/2) ^37^. Then, pool capacity, ph, water temperature and pool height were used as fixed factors in a glm with either water Hg or sediment Hg as response variables. Because there can be (unmeasured) pool characteristics that vary across pool types, we did a second set of analysis with waterHg and sedimentHg as response variables and pool type as fixed factor with six levels (canopy pool, rockpool, palm bract, treehole, tree buttress, fallen log). We accounted for possible collinearity by calculating the models’ Variance Inflation Factor, which was considered acceptable if it was < 3^51^.

#### Differences in tadpole Hg burdens across the vertical axis of the forest and in function of life history, developmental stage and environmental Hg concentrations

The five studied species were grouped according to their life history and reproductive ecology into three categories: “slow, tadpole transport” (*D. tinctorius*, *A. femoralis*), “slow, egg-eaters” (*T. resinifictrix*, *O. oophagus*), and “fast, no parental care” (*R. castaneotica*). To test our predictions that tissueHg is higher (1) at ground level than at higher altitudes from the ground; (2) in species with slow development; and (3) at the most advanced developmental stages, we ran a linear mixed model with tissueHg as a response variable and pool height, tadpole developmental stage (early, late, metamorph) and life history (fast, slow/egg-eater, slow/tadpole transport) as predictors (fixed factors) with waterHg and tadpole length as covariates and pool ID as a random factor. To account for among-species differences in size at different developmental stages, we included the interaction between tadpole length and developmental stage in our models.

#### DNA damage in relation to tadpole Hg burdens across the vertical axis of the forest and in function of life history, developmental stage and environmental Hg concentrations

To examine the influence of Hg burdens in the tadpole, as well as of reproductive ecology/life history, developmental stage and pool height on levels of DNA damage, we ran a linear mixed model with 8HOdG as a response variable and Hg burdens, tadpole height, tadpole developmental stage (early, late, metamorph) and life history (fast, slow-egg feeder, slow-tadpole transport) as predictors (fixed factors). To account for among-species differences in size at different developmental stages, we included the interaction between tadpole length and developmental stage in our models.

All continuous variables were log-transformed and centred around the mean to facilitate comparisons and reduce collinearity. We used the ‘emmeans’ function of the *emmeans* package ^52^ to perform pairwise comparisons among the categories of the variable of interest, and we applied ‘False Discovery Rate’ (FDR) correction for the p-values obtained from each model ^53^. General linear mixed models were run with the package *lme4 v.2.0.1* ^54^. Model selection was done using ∂AIC < 2 as criterion. When two models had similar support, we chose the one with the best model diagnostics. Model diagnostics were performed for all models by the ‘simulateResiduals’ function of the *DHARMa v.0.4.7* package ^55^. All statistical analyses were run in R v.4.6.0 ^56^ using the RStudio v. 2026.04.0+526 interface (Posit Team 2025).

## 3. Results

### 3.1 Effects of pool characteristics on Hg concentrations in pool water and sediment

We characterised 19 pools in 2024 and 28 pools in 2025 (height mean ± SE = 235 ± 81.7cm, range =1 – 2500 cm; N = 47). Levels of mercury in the water ranged from 0.015 to 8.540 ppm (Mean±SE = 1.620 ± 0.282; N = 42), while mercury levels in the sediment averaged 0.215 ppm and presented minimal variation (SE = 0.014, range = 0.013 – 0.494; N = 42). Therefore, in subsequent analyses we used water Hg, instead of sediment Hg, as predictor of Hg burdens in the tadpoles’ tissues. Pool water temperature had a significant effect on the amount of Hg found in the water (estimate ± SE = −11.680 ± 5.21357, t = −2.24, p = 0.031), such that pools with higher temperatures had lower amounts of Hg in the water (Fig. 1A; see Table S1 for model selection). None of the measured pool traits influenced sediment Hg (see Table S2 for model selection), but pool type did (Kruskal-Wallis rank sum test: Chi-sq = 15.449, df = 5, p = 0.009; Fig. 1B).

**Figure 1.**
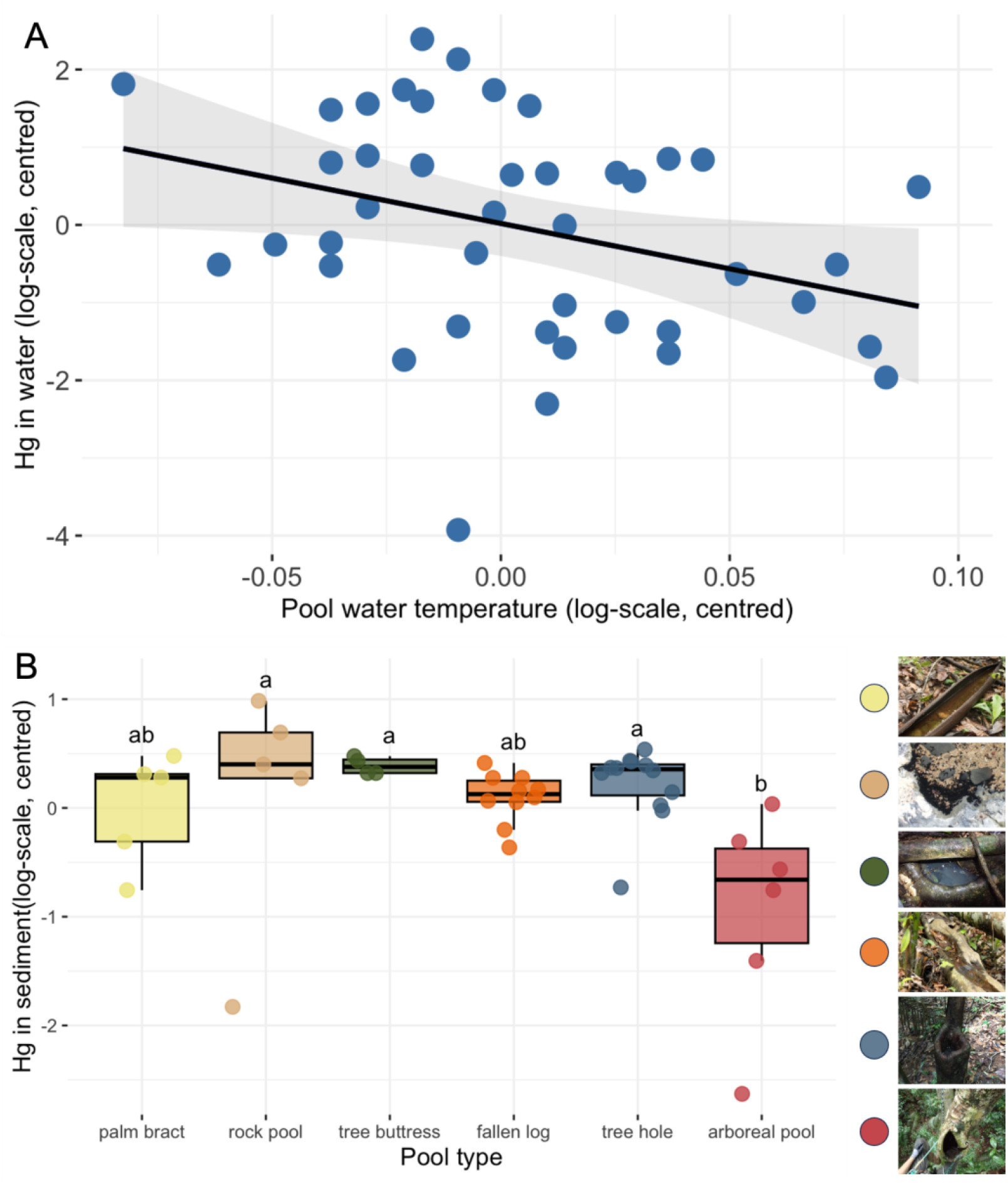
Differences in mercury (Hg) levels in (A) the water, in function of water temperature; and (B) the sediment across different pool types. Boxes show the median (horizontal line) and the 25th and 75th percentiles of data distribution and circles show individual observations.

Following a Dunn’s Test for pairwise comparisons with FDR correction, we found that sediment in canopy pools have lower Hg levels than that of rockpools (estimate = −2.88, p = 0.02), buttresses (estimate = −3.23, p =0.012) and treeholes (estimate = −3.17, p = 0.012).

### 3.2 Effects of pool height, developmental stage and reproductive ecology on Hg burdens in tadpole tissue

We measured Hg burdens in 26 tadpoles of *R. castaneotica* (between two and five individuals from nine different pools; mean±SE = 0.204 ± 0.021; range = 0.009 – 0.401 µg/g), 54 of *A. femoralis* (between one and five individuals from 16 pools; mean±SE = 0.361 ± 0.041; range = 0.019 – 1.200 µg/g), 56 of *D. tinctorius* (between one and five individuals from 30 pools; mean±SE = 0.277 ± 0.031; range = 0.011 – 1.290 µg/g), 16 of *O. oophagus* (between two and five individuals from 4 pools; mean±SE = 0.164 ± 0.053; range = 0.032 – 0.748 µg/g), and 10 of *T. resinifictrix* (five individuals two different pools; mean±SE = 0.282 ± 0.085; range = 0.020 – 0.690 µg/g). Across all species, we obtained Hg burdens for 89 tadpoles at early developmental stages (mean±SE = 0.266 ± 0.026; range = 0.011 – 1.20 µg/g), 30 at late developmental stages (mean±SE =0.266 ± 0.041; 0.009 – 0.799 µg/g), and 41 at the prometamorphic/metamorphic stage (mean±SE = 0.336 ± 0.041; range = 0.021 – 1.290 µg/g).

The best model to explain Hg burdens in the tadpoles included reproductive ecology/life history, pool height and the interaction between developmental stage and tadpole length. WaterHg was not informative and was thus excluded from the model (see Table S3 for model selection). Residual diagnostics run with DHARMA detected deviations in residual quantiles, but all other basic assumptions about overdispersion, normality, zero inflation, and outliers were met. Therefore, and given that the interaction term was biologically meaningful, we retained this model based on selection criteria (lowest AIC).

As predicted, pool height had a strong effect on the amount of Hg in the tadpoles’ tissue, with tadpoles from higher pools exhibiting significantly lower Hg burdens than their counterparts in ground pools (Table 1; Fig. 2).

**Figure 2.**
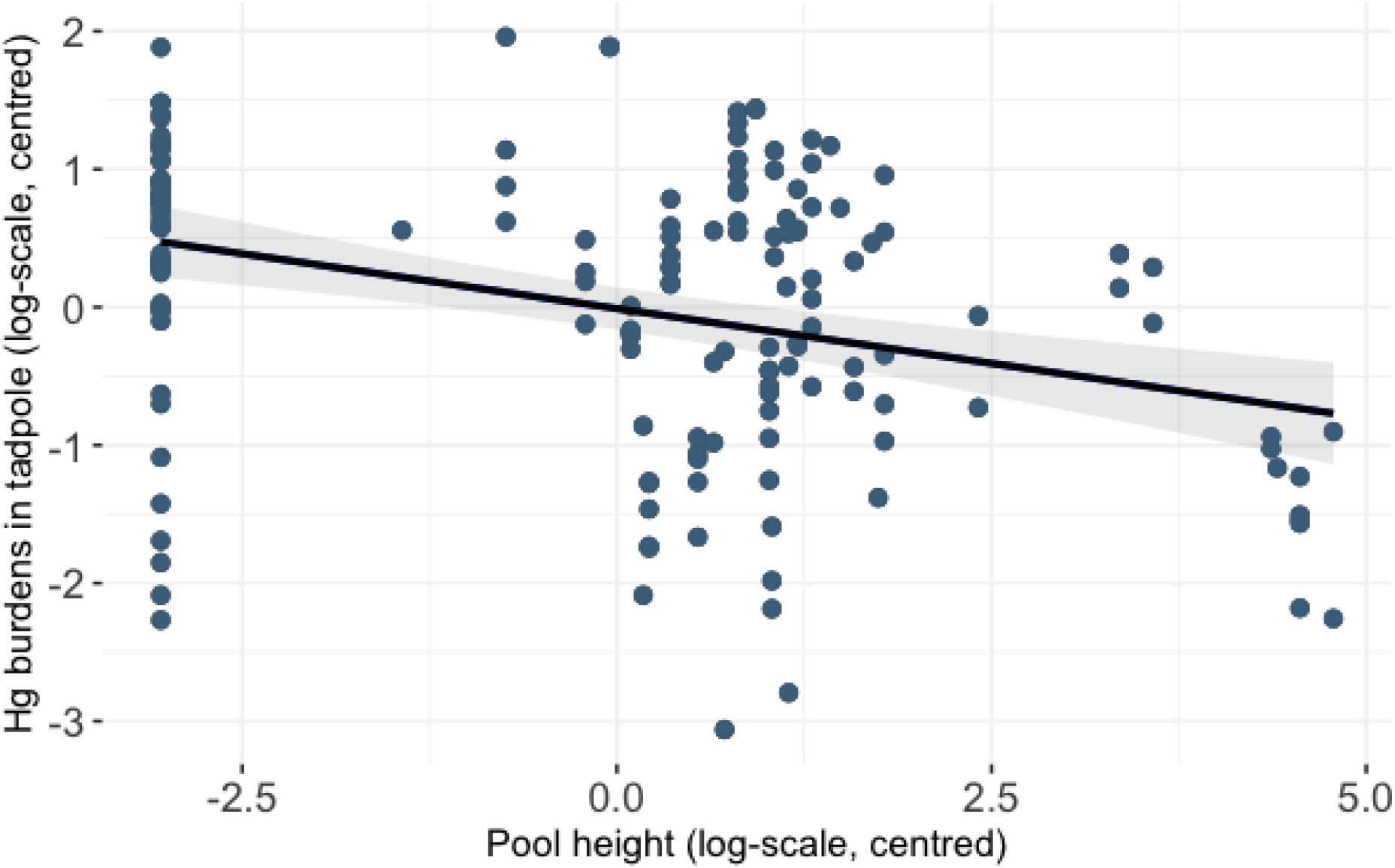
Mercury (Hg) burdens in tadpole tissue in relation to pool height. Hg burdens in tadpole tissue decrease as pool height increases.

**Table 1.**
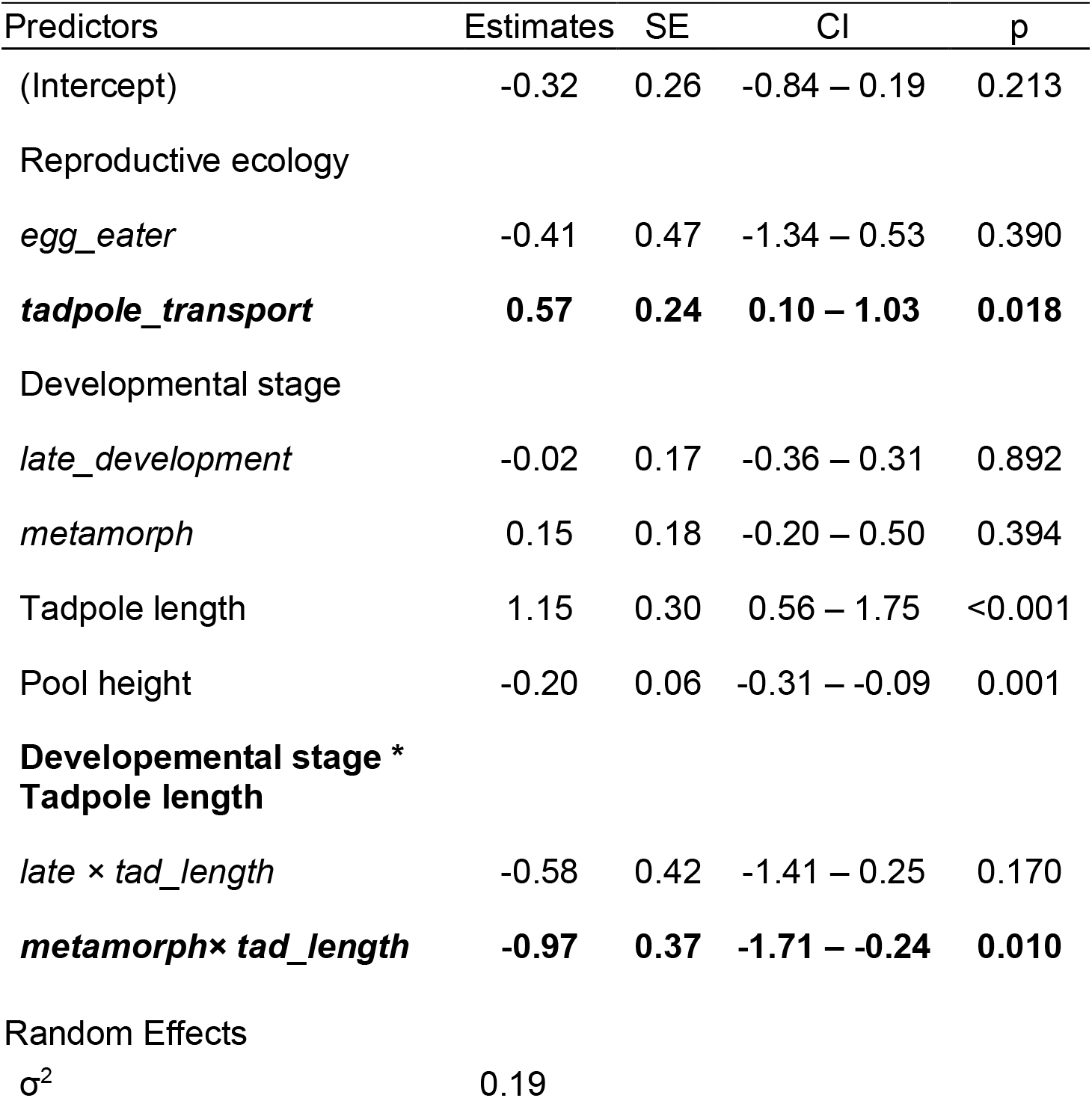
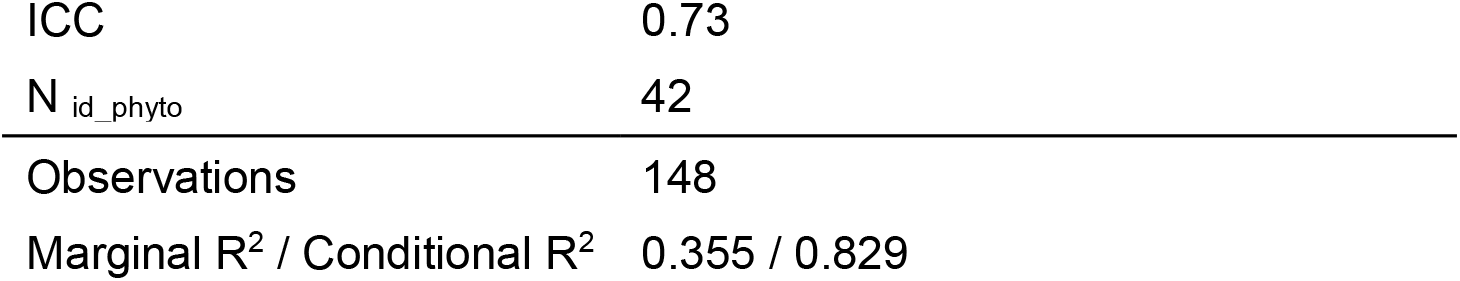
Estimates, standard error (SE), lower and upper 95 % confidence intervals (CI), and *p*-values of explanatory variables for the analysis of Hg burdens in tadpole tissue. Bold font indicates parameters with significant effects on the response variable at alpha = 0.05 and whose 95 % confidence intervals do not overlap with zero.

| Predictors | Estimates | SE | CI | p |
| --- | --- | --- | --- | --- |
| (Intercept) | -0.32 | 0.26 | -0.84 – 0.19 | 0.213 |
| Reproductive ecology |  |  |  |  |
| *egg_eater* | -0.41 | 0.47 | -1.34 – 0.53 | 0.390 |
| ***tadpole_transport*** | **0.57** | **0.24** | **0.10 – 1.03** | **0.018** |
| Developmental stage |  |  |  |  |
| *late_development* | -0.02 | 0.17 | -0.36 – 0.31 | 0.892 |
| *metamorph* | 0.15 | 0.18 | -0.20 – 0.50 | 0.394 |
| Tadpole length | 1.15 | 0.30 | 0.56 – 1.75 | <0.001 |
| Pool height | -0.20 | 0.06 | -0.31 – -0.09 | 0.001 |
| **Developmental stage \***<br>**Tadpole length** |  |  |  |  |
| *late × tad_length* | -0.58 | 0.42 | -1.41 – 0.25 | 0.170 |
| ***metamorph × tad_length*** | **-0.97** | **0.37** | **-1.71 – -0.24** | **0.010** |
| Random Effects |  |  |  |  |
| $\sigma^2$ | 0.19 | | | |

|  |  |
| --- | --- |
| ICC | 0.73 |
| N<sub>id_phyto</sub> | 42 |
| Observations | 148 |
| Marginal R<sup>2</sup> / Conditional R<sup>2</sup> | 0.355 / 0.829 |

Also, in line with our expectations, tadpoles from species with slow larval development and parental care in the form of tadpole transport (*A. femoralis*, *D. tinctorius*) had ∼76% higher Hg burdens than those with fast larval development and no parental care (*R. castaneotica*) (Fig. 3), whereas no differences were found in the Hg burdens of tadpoles with fast larval development and no parental care (*R. castaneotica*) and tadpoles with slow development and oophagy. Post hoc pairwise comparisons corrected for false discovery rate (FDR) confirmed significant differences between species with fast development/no parental care and the species with slow development/tadpole transport (estimate±SE = −0.566±0.239, df = 133.9, t ratio = −2.373, p = 0.037), and revealed that species with slow development/oophagy have also significantly lower Hg burdens than those with slow development/tadpole transport (estimate±SE = −0.973 ±0.414, df = 35.7, t ratio = −2.348, p=0.037).

**Figure 3.**
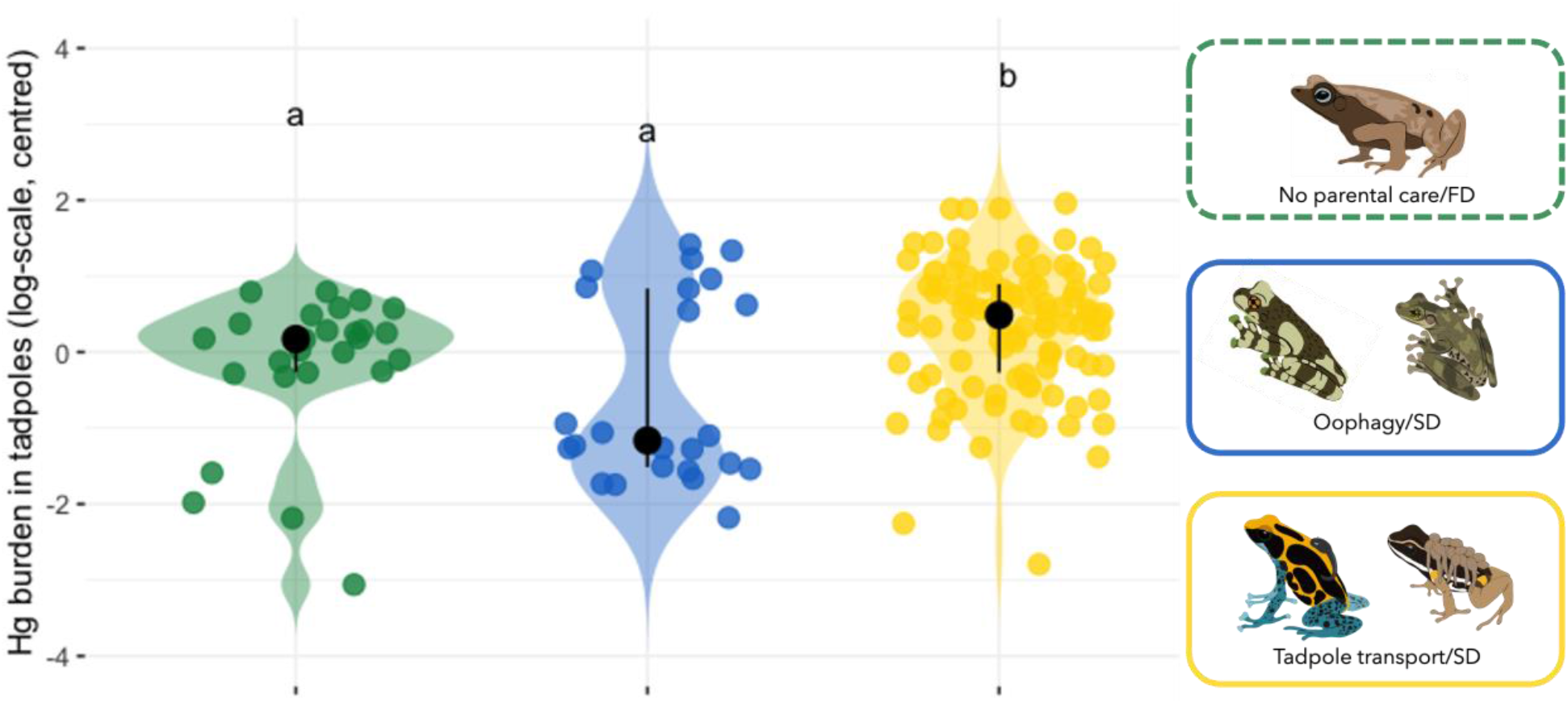
Mercury (Hg) burdens in tadpole tissue in relation to reproductive ecology/life history. Hg burdens are significantly higher in species with slow development and tadpole transport than in tadpoles with slow development and oophagy, or fast development and no parental care. Violin plots show the distribution and density of observations for each category of Reproductive Ecology. Points represent individual observations. Black points indicate the median, and vertical bars represent the central 50% interval (25th– 75th percentiles).

Differences in Hg burdens across developmental stages depended on tadpole length, such that there was a positive association between Hg burden and tadpole length in early tadpoles but not in tadpoles in the other two categories. Post hoc comparisons (FDR corrected) of the slopes showed that the relationship between tadpole length and Hg burdens has a significantly steeper slope at early developmental stages than at the metamorph stage (estimate±SE = 0.974±0.375, df = 124, t ratio = 2.598, p = 0.028), whereas no differences in slopes were found for early vs. late tadpoles (estimate±SE = 0.578±0.422, df = 121, t ratio = 1.372, p = 0.359), or late tadpoles vs. metamorphs (estimate±SE = 0.395±0.385, df = 122, t ratio = 1.026, p = 0.562) (Fig. S1).

### 3.3 Effects of Hg burdens, life history and developmental stage on DNA damage

We were able to quantify oxidative DNA damage (measured as amount of 8-Hydroxy-2′-deoxyguanosine in pg) in 26 tadpoles of *R. castaneotica* (mean±SE = 2.41 ± 0.597; range = 0.0965 – 15.9 pg), 48 of *A. femoralis* (mean±SE = 4.78 ± 1.13; range = 0.138 – 54.0), 49 of *D. tinctorius* (mean±SE = 6.02 ± 1.07; range = 0.467 – 47.7 pg), 11 of *O. oophagus* (mean±SE = 3.08 ± 0.869; range = 0.161 – 9.69 pg), and 10 of *T. resinifictrix* (mean±SE = 6.15 ± 1.49; range = 1.52 – 14.1 pg). 8-Hydroxy-2′-deoxyguanosine was below detectable levels in six individuals of *R. castaneotica,* six of *A. femoralis,* seven of *D. tinctorius,* and five of *O. oophagus*.

The best model to explain levels of oxidative DNA damage included only developmental stage and tadpole length. Reproductive ecology, pool height, Hg burdens in tadpole tissue and the interaction between developmental stage and tadpole length were excluded from the model (see Table S4 for model selection). Metamorphs had levels of DNA damage approximately 48% lower than tadpoles at the early stage (Table 2; Fig. 4A). This difference was corroborated with post hoc pairwise comparisons with FDR correction (estimate ± SE = 0.651 ± 0.214, df = 99.4, t ratio = 3.035, p = 0.009). No differences in oxidative DNA damage were found between tadpoles at late developmental stage and metamorphs (estimate ± SE = 0.332 ± 0.253, df = 132.2, t ratio = 1.310, p =0.192). Overall, larger tadpoles had higher levels of DNA damage (estimate±SE = 0.919 ± 0.240, df=95.0425, t = 3.828, p < 0.001) (Fig. 4B).

**Figure 4.**
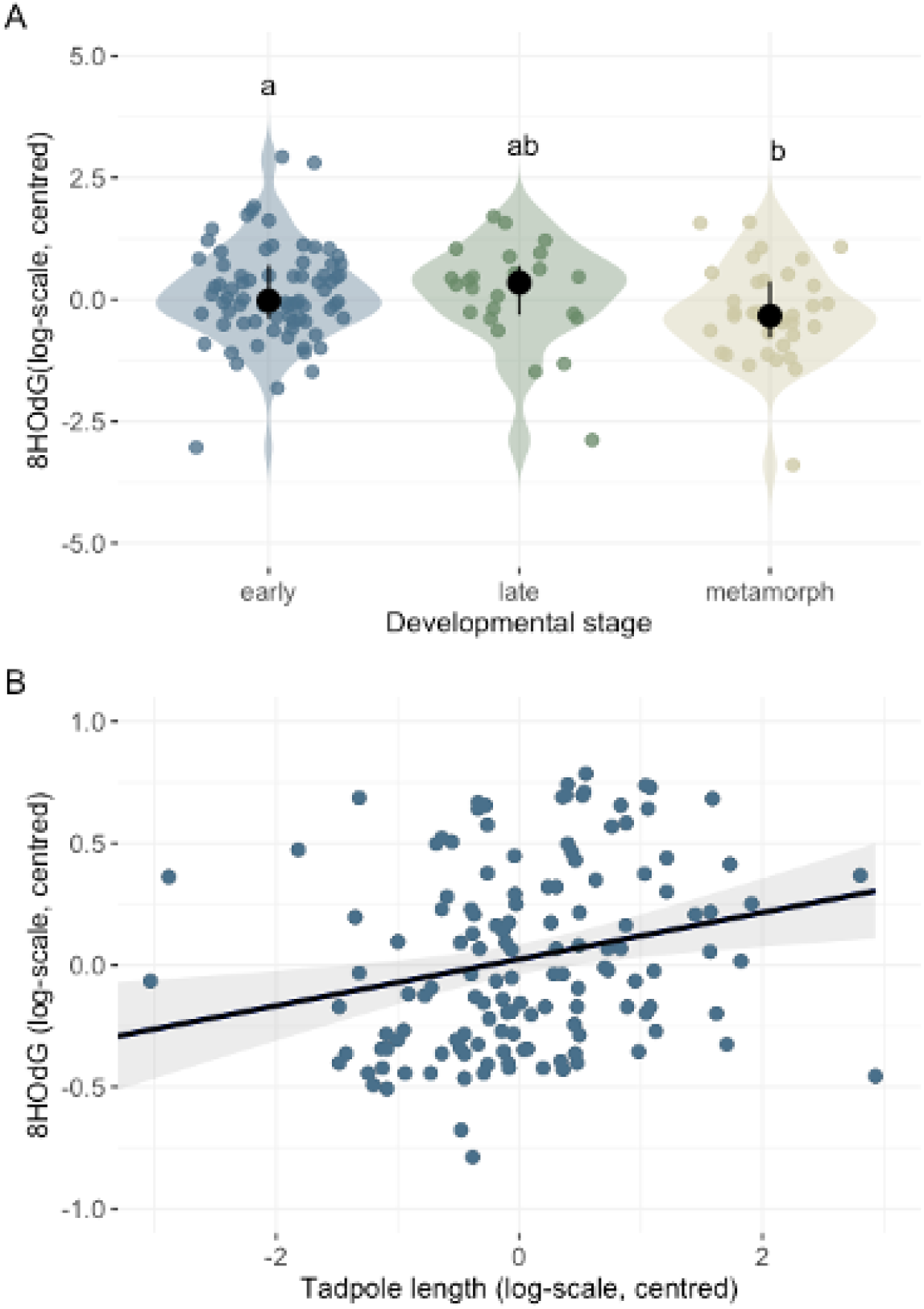
Oxidative DNA damage (measured as amount of 8HOdG) levels in relation to (A) tadpole developmental stage; and (B) tadpole length. DNA damage levels are significantly higher in tadpoles at early developmental stages than in metamorphs or tadpoles close to the metamorphic climax, and in tadpoles with larger body size. Violin plots show the distribution and density of observations for each category of Developmental Stage. Points represent individual observations. Black points indicate the median, and vertical bars represent the central 50% interval (25th–75th percentiles).

**Table 2.** Estimates, standard error (SE), lower and upper 95 % confidence intervals (CI), and *p*-values of explanatory variables for the analysis of levels of DNA damage across tadpoles. Bold font indicates parameters with significant effects on the response variable at alpha = 0.05 and whose 95 % confidence intervals do not overlap with zero.

| Predictors | Estimates | SE | CI | p |
| --- | --- | --- | --- | --- |
| (Intercept) | 0.22 | 0.12 | -0.01 – 0.45 | 0.064 |
| Developmental stage |  |  |  |  |
| *late_development* | -0.32 | 0.23 | -0.78 – 0.14 | 0.172 |
| ***metamorph*** | **-0.65** | **0.21** | **-1.06 – -0.24** | **0.002** |
| **Tadpole length** | **0.92** | **0.24** | **0.44 – 1.39** | **<0.001** |
| **Random Effects** |  |  |  |  |
| $\sigma^2$ | 0.76 | | | |
| ICC | 0.12 |  |  |  |
| N<sub>id_phyto</sub> | 47 |  |  |  |
| Observations | 137 |  |  |  |
| Marginal R<sup>2</sup> / Conditional R<sup>2</sup> | 0.137 / 0.245 |  |  |  |

## Discussion

In this study, we present the first comprehensive multi-species assessment of Hg contamination, its ecological correlates, and its consequences on DNA integrity in phytotelm-breeding amphibians. Our findings highlight the overlooked importance of phytotelmata as Hg repositories and sources, even in a site relatively isolated from anthropogenic activities. Furthermore, we reveal a vertical gradient in Hg contamination and a differential Hg accumulation in relation to life history strategies and reproductive ecology.

### Predictors of environmental and tissue Hg concentrations

Neither pool height, nor pool capacity had an influence on the amount of Hg found in the water or sediment. However, increased water temperature was related to lower levels of Hg in the water (see Fig. 1A. This pattern could be explained either by volatilisation of inorganic Hg or Hg methylation, both of which are known to increase with increasing temperatures^13^, ^57^. It is thus possible that warmer pools with low total Hg concentrations have a higher proportion of MeHg relative to inorganic Hg, a pattern we cannot detect with our measurements. The lack of effect of pH on water and sediment Hg is unexpected, as it has been found to influence environmental concentrations of Hg^13^, ^58^. The type of phytotelmata was not related to the Hg concentration in the water, but canopy pools had clearly lower Hg concentrations in the sediment than almost any other pool type. One possible explanation for these findings is the time of the year at which Hg contamination was measured. High levels of rainfall might contribute to the dilution of Hg concentrations and pH variability in phytotelmata and other bodies of water. This is particularly true for canopy pools, as treeholes in the forest canopy are more exposed to rainfall input due to higher levels of storage and interception compared to lower levels of the forest ^59^.

Contrary to our expectations, and to previous studies^60^, ^61^, water Hg was not a good predictor of Hg burdens in the tadpoles. This could also be explained by the fact that the measurements were done during the rainy season, as explained above. While Hg burdens in the tadpoles reflect bioaccumulation over time, our measurements of water Hg are likely reflecting only a “snapshot” of the dynamics of Hg in these pools. Future measurements during the dry season may provide a clearer picture of how Hg concentrations fluctuate across seasons, particularly in the sediment, as many pools are likely to dry out almost completely during the dry period. In addition, we cannot discard that unmeasured pool characteristics, such as dissolved oxygen concentration, influence the observed Hg levels in the water, as has been previously shown ^13^, ^60^.

In line with our predictions, there was a vertical gradient in Hg burdens, with tadpoles from canopy pools exhibiting significantly lower burdens compared to tadpoles from pools on the ground. This contamination gradient might reflect the accumulation of mercury due to increased deposition from litterfall after being deposited in canopy foliage 4. Higher amounts of organic matter and fine sediment in ground pools can create low-oxygen conditions that favour the microbial conversion of inorganic mercury into methylmercury ^13^, ^62^. The bacterial and fungal biofilms which grow under these circumstances might in turn contaminate the sediments and leaves on which tadpoles will graze, thus contributing to increased Hg uptake ^61^, as has been suggested in wood frog tadpoles ^63^. By contrast, high arboreal pools are often more exposed to sunlight and UV radiation, a combination thought to reduce methylmercury production ^13^ and which could explain why tadpoles from high arboreal pools had lower mercury concentrations.

As Hg is bioaccumulated over time, tadpoles of species with a longer larval phase are expected to accumulate more Hg in their tissues before metamorphosis. In line with this, the species with the shortest larval period in this study, *R. castaneotica*, exhibited the lowest Hg burdens. Their rapid development likely limits the time available for environmental exposure and diet-related bioaccumulation, reducing mercury burdens at the tadpole stage. This pattern has been previously shown for other heavy metals ^64^, even in habitats with elevated contamination. Our findings, however, show that having a slow development alone does not explain higher bioaccumulation. In our study, four of the five species involved had a relatively slow development, but differed in other aspects of their ecology. Two species exhibit either facultative or obligatory oophagy, with their larval phase occurring almost exclusively in arboreal pools. The other two, which had the highest Hg burdens, are omnivorous, feeding mostly on detritus, and are transported to phytotelmata as tadpoles by their parents. One of these species, *A. femoralis*, occupies exclusively ground pools, whereas the other one, *D. tinctorius*, uses pools all the way from the ground to the canopy ^9^. It is known that Hg amounts in the forest floor are higher than in the canopy due, among other factors, to litterfall and throughfall. For example, in some boreal forests, it is estimated that the flux of Hg in throughfall and litterfall under the canopy can be 2-3 times greater than the flux by direct wet deposition ^65^. Therefore, the detritus these two species forage on is likely more contaminated over time than that in the canopy pools. In addition to this, *D. tinctorius* is known to be carnivorous and an opportunistic cannibal ^37^, ^66^. Given that Hg biomagnifies across the food web ^14^, ^67^, it is likely that feeding on already contaminated con- or heterospecifics increases Hg uptake in *D. tinctorius* tadpoles. Interestingly, however, *A. femoralis* and *D. tinctorius* did not differ in Hg burdens. In fact, the Hg burdens of *D. tinctorius* tadpoles from ground pools were approximately 21% lower than those of *A. femoralis* tadpoles (see Fig S2), suggesting that cannibalism is not the main reason for the high levels of Hg found in these two species. Instead, grazing on contaminated detritus during their long larval phase appears to be the most plausible explanation for the high Hg burdens in their bodies.

Despite their prolonged larval development, tadpoles of species with either obligatory or facultative oophagy had significantly lower Hg burdens than tadpole transporters, which may point at efficient detoxification and/or elimination mechanisms. Previous research suggests that at low and medium exposure levels, as those in arboreal pools, Hg elimination may hinder toxicity in tadpoles ^68^. These mechanisms, however, may not be sufficient to depurate Hg at high exposure levels ^68^. An alternative, non-mutually exclusive explanation, is that maternal transfer may be particularly low in these arboreal species. Despite mounting evidence pointing at maternal transfer as an established Hg exposure pathway for offspring in multiple taxa ^69^, ^70^, work on some arthropods, birds and mammals has revealed that females might be capable of modulating the amount of Hg they pass on to their offspring ^71^, particularly when the female has high Hg burdens herself ^70^, ^72^. Furthermore, the maternal transfer of Hg varies in magnitude across species^70^, which could also contribute to explaining our findings. In this study we did not measure the Hg burdens in adult frogs or explored detoxification or elimination mechanisms, and are therefore unable to provide further support to either explanation.

### DNA damage and Hg burdens

Contrary to our predictions, we found no effect of Hg burdens on the levels of oxidative DNA damage observed in tadpoles, which is surprising considering the known deleterious effects that Hg can exert on vital macromolecules ^73^. One possible explanation is that such damages are only inflicted above a certain threshold of Hg burdens, far above the ones observed in the tadpoles in this study. However, metamorphs had overall lower levels of DNA damage than tadpoles at earlier developmental stages. This pattern may reflect the physiological and cellular remodelling that occurs during metamorphosis, which involves enhanced DNA repair and turnover of damaged cells. Similar to what occurs in insects^74^, amphibian metamorphosis involves programmed cell death and proliferation and differentiation of *de novo*, physiologically younger cells ^75^, which could help mitigate the detrimental effects of prior early-life stress exposure. With the present results, however, we cannot discard that the low levels of DNA damage found in metamorphs are the byproduct of selective mortality, in which case, individuals with high levels of DNA damage at earlier stages may not have survived to the onset of metamorphosis. Although direct evidence for such a process in amphibians is lacking, high 8-OHdG levels have been linked with subsequent mortality in other vertebrate species, including humans ^76^. Future studies employing longitudinal sampling across developmental stages will be necessary to determine the true within-individual pattern of age-related variation in oxidative DNA damage and to distinguish it from changes arising through selective mortality.

Taken together, our findings demonstrate that both environmental and ecological traits can shape mercury exposure and accumulation in tropical amphibians and underscore the importance of integrating environmental and biological perspectives in future ecotoxicological assessments in the wild in changing tropical landscapes.

## Acknowledgements

We are most grateful to Floriana Stanca for help in field data collection; Rosanna Mangione for doing the Hg quantification in some of the samples; Clara Ferreira for help with DNA damage estimation; Sara Calhim for statistical advice; and Susanne Stückler for valuable comments on an early version of this MS. We are indebted to the Nouragues Ecological Research Station (managed by CNRS), which benefits from “Investissement d’Avenir” grants managed by Agence Nationale de la Recherche (AnaEE France ANR-11-INBS-0001; Labex CEBA ANR-10-LABX-25-01), for logistic support. BR and AP are part of a multi-researcher partnership with the Nouragues Nature Reserve (Convention Cadre de Partenariat N◦01-2019) aimed at improving and spreading knowledge about amphibians.

## Author contributions

**Bibiana Rojas:** conceptualization (equal), methodology (equal), data curation (equal), formal analysis (lead), visualization (lead), writing – original draft (lead), writing – review and editing (equal), funding acquisition (equal), resources (lead), project administration (equal). **Paul Kesseler:** methodology (supporting), investigation (equal), data curation (equal), writing – original draft (supporting), writing – review and editing (equal). **Lia Schlippe Justicia:** methodology (supporting), investigation (equal), data curation (equal), writing – original draft (supporting), writing – review and editing (equal). **Andrius Pašukonis:** methodology (supporting), investigation (supporting), writing – review and editing (equal). **Joan Espigol:** investigation (supporting), writing – review and editing (equal). **Doriane Hagnier:** investigation (supporting), writing – review and editing (equal). **Shubham Soni:** investigation (supporting), writing – review and editing (equal). **Valentine Alt:** methodology (supporting), investigation (supporting), writing – review and editing (equal). **Jérémy Lemaire:** conceptualization (equal), methodology (equal), mercury quantification (lead), writing – original draft (supporting), writing – review and editing (equal), resources (supporting). **Valeria Marasco:** conceptualization (equal), methodology (equal), DNA damage quantification (lead), writing – original draft (supporting), writing – review and editing (equal), resources (supporting), funding acquisition (equal), resources (supporting), project administration (equal).

---

**Table S1.** WaterHg Model Selection. 1.1 WaterHgmodel, AIC = 151.7873

| Characteristic | Beta | 95% CI | p-value |
| --- | --- | --- | --- |
| centlogcapacity | 0.07 | -0.15, 0.30 | 0.5 |
| centlogwatertemp | -12 | -23, -0.76 | **0.043** |
| ph | 0.01 | -0.68, 0.70 | >0.9 |
| centlogheight | -0.12 | -0.31, 0.08 | 0.2 |
| Abbreviation: CI = Confidence Interval |  |  |  |

**1.2.** WaterHgmodel2, AIC = 149.7886

**1.2 WaterHgmodel2, AIC = 149.7886**
| Characteristic | Beta | 95% CI | p-value |
| --- | --- | --- | --- |
| centlogcapacity | 0.07 | -0.15, 0.29 | 0.5 |
| centlogwatertemp | -12 | -22, -1.5 | **0.031** |
| centlogheight | -0.12 | -0.31, 0.07 | 0.2 |
| Abbreviation: CI = Confidence Interval |  |  |  |

**1.3.** WaterHgmodel3, AIC = 148.2459

**1.3 WaterHgmodel3, AIC = 148.2459**
| Characteristic | Beta | 95% CI | p-value |
| --- | --- | --- | --- |
| centlogwatertemp | -12 | -22, -1.3 | **0.032** |
| centlogheight | -0.10 | -0.28, 0.08 | 0.3 |
| Abbreviation: CI = Confidence Interval |  |  |  |

**1.4.** WaterHgmodel4, AIC = 147.4465

**1.4 WaterHgmodel4, AIC = 147.4465**
| Characteristic | Beta | 95% CI | p-value |
| --- | --- | --- | --- |
| centlogwatertemp | -12 | -22, -1.5 | **0.031** |
| Abbreviation: CI = Confidence Interval |  |  |  |

**Table S2.** SedimentHg Model Selection. 2.1 SedimentHgmodel 1, AIC = 28.28108

| **Characteristic** | **Beta** | **95% CI** | **p-value** |
| --- | --- | --- | --- |
| centlogcapacity | 0.03 | -0.31, 0.38 | 0.9 |
| centlogwatertemp | -1.2 | -13, 10 | 0.9 |
| ph | -0.02 | -0.84, 0.79 | >0.9 |
| centlogheight | 0.02 | -0.12, 0.17 | 0.8 |
| Abbreviation: CI = Confidence Interval |  |  |  |

**2.2.** SedimentHgmodel 2, AIC = 26.28701

2.2. SedimentHgmodel 2, AIC = 26.28701
| **Characteristic** | **Beta** | **95% CI** | **p-value** |
| --- | --- | --- | --- |
| centlogcapacity | 0.03 | -0.29, 0.35 | 0.8 |
| centlogwatertemp | -1.3 | -11, 8.0 | 0.8 |
| centlogheight | 0.02 | -0.11, 0.15 | 0.7 |
| Abbreviation: CI = Confidence Interval |  |  |  |

**2.3.**
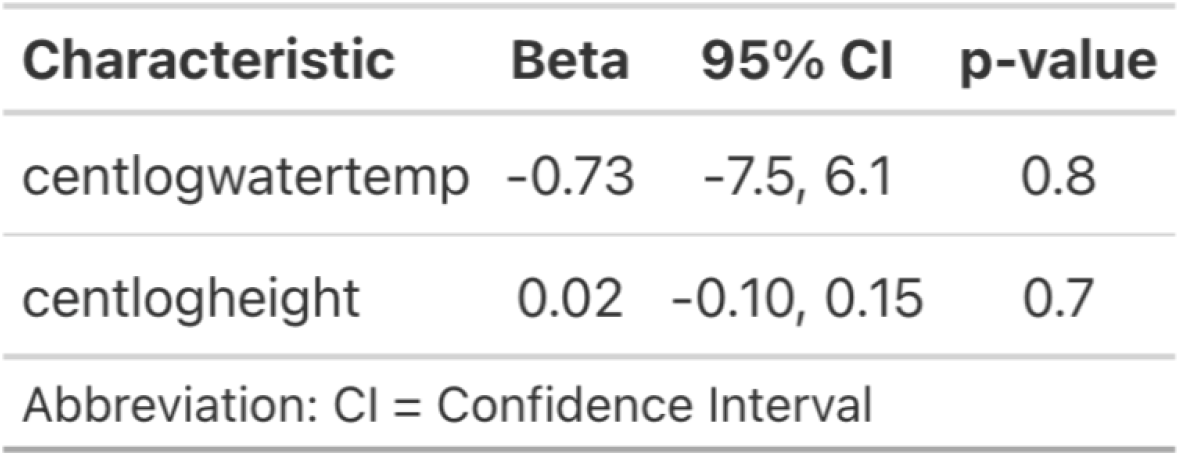
SedimentHgmodel 3, AIC = 24.35215

**2.4.** SedimentHgmodel 4, AIC = 22.41277

2.4 SedimentHgmodel 4, AIC = 22.41277
| Characteristic | Beta | 95% CI | p-value |
| --- | --- | --- | --- |
| centlogheight | 0.02 | -0.09, 0.13 | 0.7 |
| Abbreviation: CI = Confidence Interval |  |  |  |

**Table S3.** Model Selection Hg burdens in tadpole tissue. 3.1 Full model, AIC = 298.4749

| Characteristic | Beta | 95% CI | p-value |
| --- | --- | --- | --- |
| centlogwaterhg | 0.13 | -0.05, 0.31 | 0.2 |
| rep_strat |  |  |  |
| no_care | — | — |  |
| egg_eater | -0.46 | -1.4, 0.48 | 0.3 |
| tad_trans | 0.56 | 0.09, 1.0 | **0.019** |
| tad_stage_cat |  |  |  |
| early | — | — |  |
| late | -0.04 | -0.38, 0.29 | 0.8 |
| metamorph | 0.14 | -0.22, 0.49 | 0.4 |
| centlogtadlength | 1.1 | 0.51, 1.7 | **<0.001** |
| centlogheight | -0.18 | -0.29, -0.07 | **0.002** |
| tad_stage_cat * centlogtadlength |  |  |  |
| late * centlogtadlength | -0.49 | -1.3, 0.35 | 0.2 |
| metamorph * centlogtadlength | -0.92 | -1.7, -0.18 | **0.015** |

Abbreviation: CI = Confidence Interval

**3.2.** HgBurdens model 2, AIC = 295.56

3.2 HgBurdens model 2, AIC = 295.56
| Characteristic | Beta | 95% CI | p-value |
| --- | --- | --- | --- |
| rep_strat |  |  |  |
| no_care | — | — |  |
| egg_eater | -0.41 | -1.4, 0.54 | 0.4 |
| tad_trans | 0.57 | 0.10, 1.0 | **0.018** |
| tad_stage_cat |  |  |  |
| early | — | — |  |
| late | -0.02 | -0.36, 0.31 | 0.9 |
| metamorph | 0.15 | -0.20, 0.51 | 0.4 |
| centlogtadlength | 1.2 | 0.56, 1.8 | **<0.001** |
| centlogheight | -0.20 | -0.31, -0.08 | **0.001** |
| tad_stage_cat * centlogtadlength |  |  |  |
| late * centlogtadlength | -0.58 | -1.4, 0.25 | 0.2 |
| metamorph * centlogtadlength | -0.97 | -1.7, -0.24 | **0.010** |

Abbreviation: CI = Confidence Interval

**Table S4.** Model Selection DNA damage. 4.1 DNA damage model 1, AIC = 390.597

| Characteristic | Beta | 95% CI | p-value |
| --- | --- | --- | --- |
| centlogtissuehg | 0.14 | -0.08, 0.36 | 0.2 |
| centlogheight | 0.09 | 0.00, 0.19 | 0.057 |
| rep_strat |  |  |  |
| no_care | — | — |  |
| egg_eater | -0.70 | -1.6, 0.21 | 0.13 |
| tad_trans | -0.16 | -0.87, 0.55 | 0.7 |
| tad_stage_cat |  |  |  |
| early | — | — |  |
| late | -0.07 | -0.62, 0.47 | 0.8 |
| metamorph | -0.71 | -1.3, -0.17 | **0.011** |
| centlogtadlength | 0.51 | -0.33, 1.3 | 0.2 |
| tad_stage_cat * centlogtadlength |  |  |  |
| late * centlogtadlength | -0.20 | -1.6, 1.2 | 0.8 |
| metamorph * centlogtadlength | 0.93 | -0.29, 2.1 | 0.13 |

Abbreviation: CI = Confidence Interval

**4.2.** DNA damage model 2, AIC = 388.6475

4.2 DNA damage model 2, AIC = 388.6475
| Characteristic | Beta | 95% CI | p-value |
| --- | --- | --- | --- |
| centlogtissuehg | 0.16 | -0.03, 0.36 | 0.10 |
| centlogheight | 0.08 | -0.01, 0.16 | 0.078 |
| tad_stage_cat |  |  |  |
| early | — | — |  |
| late | -0.09 | -0.59, 0.41 | 0.7 |
| metamorph | -0.69 | -1.1, -0.28 | **0.001** |
| centlogtadlength | 0.52 | -0.30, 1.3 | 0.2 |
| tad_stage_cat * centlogtadlength |  |  |  |
| late * centlogtadlength | -0.51 | -1.8, 0.81 | 0.4 |
| metamorph * centlogtadlength | 0.60 | -0.44, 1.6 | 0.3 |

Abbreviation: CI = Confidence Interval

**4.3.**
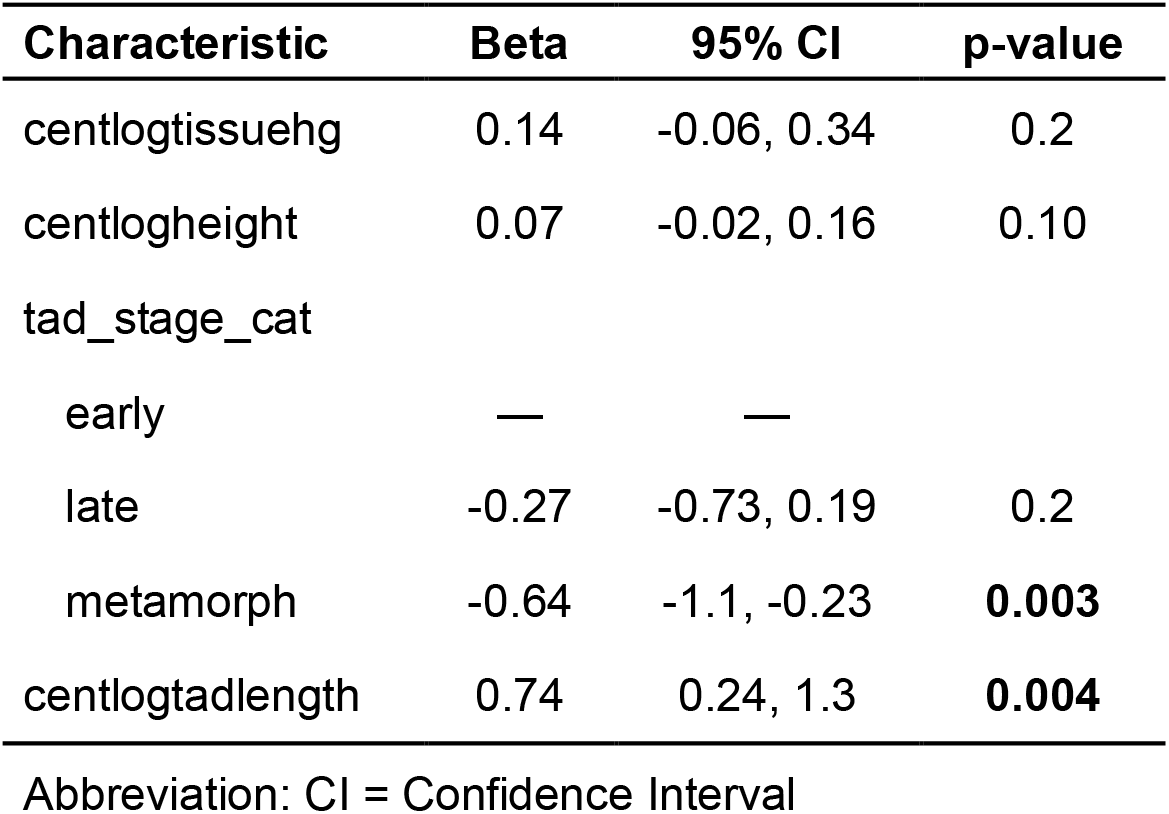
DNA damage model 3, AIC = 389.3326

**4.4.** DNA damage model 4, AIC = 386.5379

4.4 DNA damage model 4, AIC = 386.5379
| Characteristic | Beta | 95% CI | p-value |
| --- | --- | --- | --- |
| centlogheight | 0.05 | -0.03, 0.13 | 0.2 |
| tad_stage_cat |  |  |  |
| early | — | — |  |
| late | -0.31 | -0.77, 0.15 | 0.2 |
| metamorph | -0.64 | -1.1, -0.23 | **0.003** |
| centlogtadlength | 0.85 | 0.37, 1.3 | **<0.001** |

Abbreviation: CI = Confidence Interval

**4.5.**
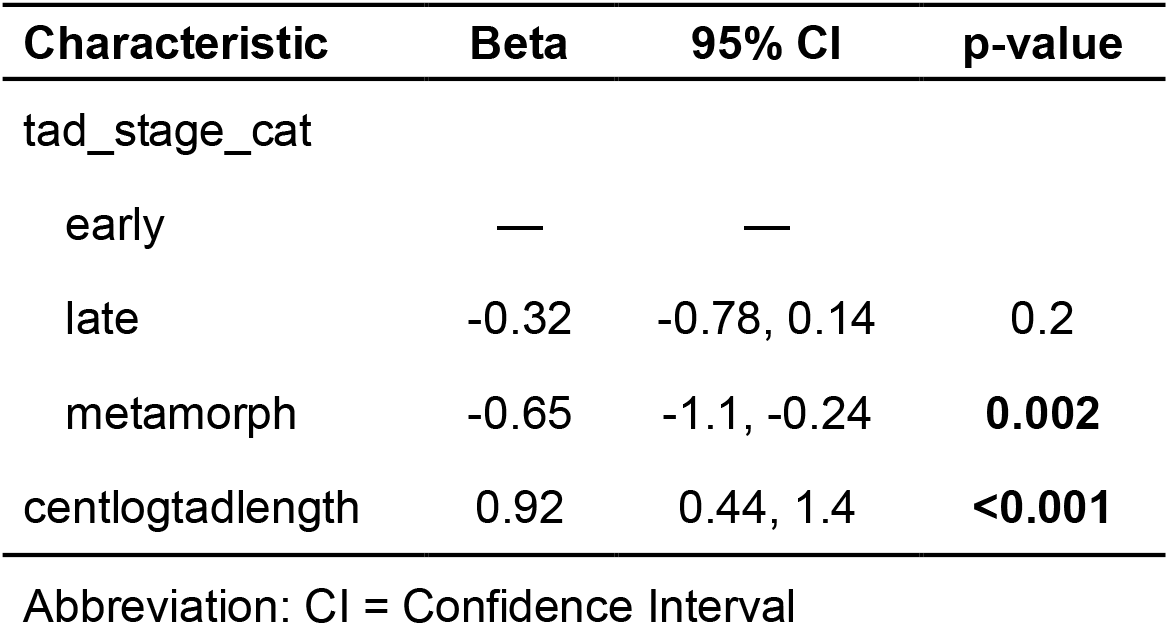
DNA damage model 5, AIC = 381.3401

4.5 DNA damage model 5, AIC = 381.3401
| Characteristic | Beta | 95% CI | p-value |
| --- | --- | --- | --- |
| tad_stage_cat |  |  |  |
| early | — | — |  |
| late | -0.32 | -0.78, 0.14 | 0.2 |
| metamorph | -0.65 | -1.1, -0.24 | **0.002** |
| centlogtadlength | 0.92 | 0.44, 1.4 | **<0.001** |

Abbreviation: CI = Confidence Interval

**Figure S1.**
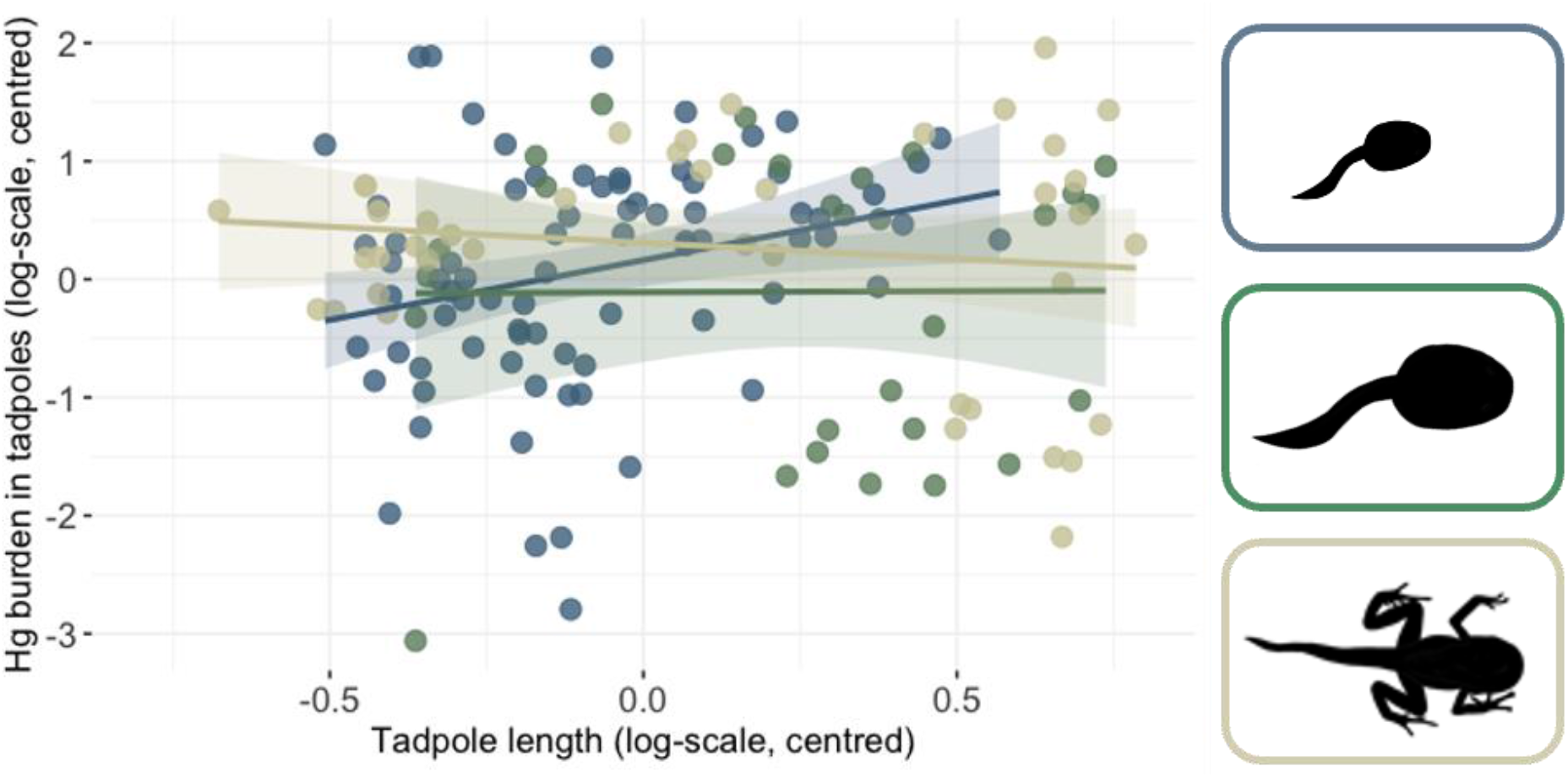
Mercury (Hg) burdens in relation to tadpole length for tadpoles at different developmental stages (i.e. early (blue) and late (green) larval developmental stages, and metamorph (kakhi)). Shaded areas show 95% confidence intervals.

**Figure S2.**
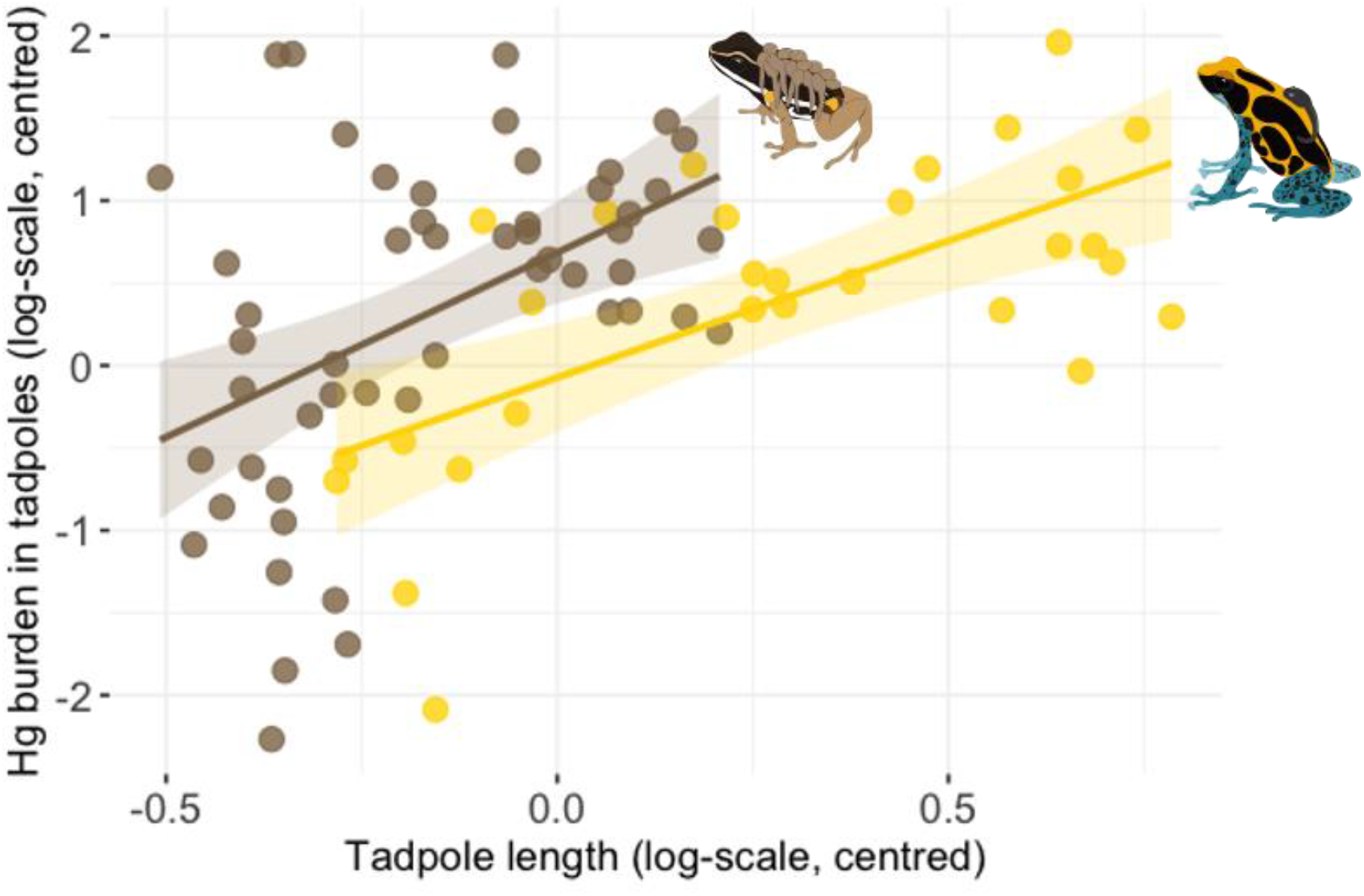
Differences in Hg burdens between *Dendrobates tinctorius* and *Allobates femoralis* tadpoles from ground pools.

## References

(1) Qiu, X.; Liu, M.; Zhang, Y.; Zhang, Q.; Lin, H.; Cai, X.; Li, J.; Dai, R.; Zheng, S.; Wang, J.;, et al. Declines in anthropogenic mercury emissions in the Global North and China offset by the Global South. Nature Communications 2025, 16 (1), 1179. DOI: 10.1038/s41467-025-56274-2.

(2) Durán, A. P.; Rauch, J.; Gaston, K. J. Global spatial coincidence between protected areas and metal mining activities. Biological Conservation 2013, 160, 272–278. DOI: 10.1016/j.biocon.2013.02.003.

(3) Gerson, J. R.; Szponar, N.; Zambrano, A. A.; Bergquist, B.; Broadbent, E.; Driscoll, C. T.; Erkenswick, G.; Evers, D. C.; Fernandez, L. E.; Hsu-Kim, H. Amazon forests capture high levels of atmospheric mercury pollution from artisanal gold mining. Nature communications 2022, 13 (1), 559.

(4) Teixeira, D. C.; Lacerda, L. D.; Silva-Filho, E. V. Mercury sequestration by rainforests: The influence of microclimate and different successional stages. Chemosphere 2017, 168, 1186–1193. DOI: 10.1016/j.chemosphere.2016.10.081.

(5) Teixeira, D. C.; Montezuma, R. C.; Oliveira, R. R.; Silva-Filho, E. V. Litterfall mercury deposition in Atlantic forest ecosystem from SE – Brazil. Environmental Pollution 2012, 164, 11–15. DOI: 10.1016/j.envpol.2011.10.032.

(6) Fostier, A. H.; Melendez-Perez, J. J.; Richter, L. Litter mercury deposition in the Amazonian rainforest. Environmental Pollution 2015, 206, 605–610. DOI: 10.1016/j.envpol.2015.08.010.

(7) Wang, X.; Bao, Z.; Lin, C.-J.; Yuan, W.; Feng, X. Assessment of global mercury deposition through litterfall. Environmental Science & Technology 2016, 50 (16), 8548–8557. DOI: 10.1021/acs.est.5b06351.

(8) Schlippe Justicia, L.; Pašukonis, A.; Rojas, B. Phytotelmata. Current Biology 2025, 35 (December 1), R1132–R1134.

(9) Fouilloux, C. A.; Serrano Rojas, S. J.; Carvajal-Castro, J. D.; Valkonen, J. K.; Gaucher, P.; Fischer, M.-T.; Pašukonis, A.; Rojas, B. Pool choice in a vertical landscape: Tadpole-rearing site flexibility in phytotelm-breeding frogs. Ecology and Evolution 2021, 11 (n/a), 9021–9038, 10.1002/ece3.7741. DOI: 10.1002/ece3.7741 (acccessed 2021/06/25).

(10) Kitching, R. L. Food webs in phytotelmata: “Bottom-Up” and “Top-Down” explanations for community structure. Annual Review of Entomology 2001, 46 (1), 729–760. DOI: 10.1146/annurev.ento.46.1.729 (acccessed 2018/06/27).

(11) Schlippe-Justicia, L.; Lemaire, J.; Dittrich, C.; Mayer, M.; Bustamante, P.; Rojas, B. Poison in the nursery: Mercury contamination in the tadpole-rearing sites of an Amazonian frog. Science of The Total Environment 2024, 912, 169450. DOI: 10.1016/j.scitotenv.2023.169450.

(12) Ma, M.; Du, H.; Wang, D. Mercury methylation by anaerobic microorganisms: A review. Critical Reviews in Environmental Science and Technology 2019, 49 (20), 1893–1936. DOI: 10.1080/10643389.2019.1594517.

(13) Ullrich, S. M.; Tanton, T. W.; Abdrashitova, S. A. Mercury in the aquatic environment: A review of factors affecting methylation. Critical Reviews in Environmental Science and Technology 2001, 31 (3), 241–293. DOI: 10.1080/20016491089226.

(14) Kidd, K.; Clayden, M.; Jardine, T. Bioaccumulation and Biomagnification of Mercury through Food Webs. In Environmental Chemistry and Toxicology of Mercury, 2011; pp 453–499.

(15) Eagles-Smith, C. A.; Silbergeld, E. K.; Basu, N.; Bustamante, P.; Diaz-Barriga, F.; Hopkins, W. A.; Kidd, K. A.; Nyland, J. F. Modulators of mercury risk to wildlife and humans in the context of rapid global change. Ambio 2018, 47 (2), 170–197. DOI: 10.1007/s13280-017-1011-x.

(16) Chételat, J.; Ackerman, J. T.; Eagles-Smith, C. A.; Hebert, C. E. Methylmercury exposure in wildlife: A review of the ecological and physiological processes affecting contaminant concentrations and their interpretation. Science of The Total Environment 2020, 711, 135117. DOI: 10.1016/j.scitotenv.2019.135117. Marasco, V.; Costantini, D. Signaling in a polluted world: oxidative stress as an overlooked mechanism linking contaminants to animal communication. Frontiers in Ecology and Evolution 2016, 4, Mini Review. DOI: 10.3389/fevo.2016.00095. Varian-Ramos, C. W.; Swaddle, J. P.; Cristol, D. A. Mercury reduces avian reproductive success and imposes selection: an experimental study with adult- or lifetime-exposure in zebra finch. PLOS ONE 2014, 9 (4), e95674. DOI: 10.1371/journal.pone.0095674. Swaddle, J. P.; Diehl, T. R.; Taylor, C. E.; Fanaee, A. S.; Benson, J. L.; Huckstep, N. R.; Cristol, D. A. Exposure to dietary mercury alters cognition and behavior of zebra finches. Current Zoology 2017, 63 (2), 213-219. DOI: 10.1093/cz/zox007 (acccessed 2/15/2023). White, A. E.; Cristol, D. A. Plumage coloration in belted kingfishers Megaceryle alcyon at a mercury-contaminated river. Waterbirds 2014, 37 (2), 144-152. DOI: 10.1675/063.037.0203. Lemaire, J.; Bustamante, P.; Mangione, R.; Marquis, O.; Churlaud, C.; Brault-Favrou, M.; Parenteau, C.; Brischoux, F. Lead, mercury, and selenium alter physiological functions in wild caimans (Caiman crocodilus). Environmental Pollution 2021, 286, 117549. DOI: doi.org/10.1016/j.envpol.2021.117549.

(17) Farina, M.; Aschner, M. Methylmercury-induced neurotoxicity: focus on pro-oxidative events and related consequences. In Neurotoxicity of Metals, Aschner, M., Costa, L. G. Eds.; Springer International Publishing, 2017; pp 267–286. Teixeira, F. B.; de Oliveira, A. C. A.; Leão, L. K. R.; Fagundes, N. C. F.; Fernandes, R. M.; Fernandes, L. M. P.; da Silva, M. C. F.; Amado, L. L.; Sagica, F. E. S.; de Oliveira, E. H. C.; et al. Exposure to inorganic mercury causes oxidative stress, cell death, and functional deficits in the motor cortex. Frontiers in Molecular Neuroscience 2018, Volume 11 - 2018, Original Research. DOI: 10.3389/fnmol.2018.00125.

(18) Martindale, J. L.; Holbrook, N. J. Cellular response to oxidative stress: Signaling for suicide and survival. Journal of Cellular Physiology 2002, 192 (1), 1–15. DOI: 10.1002/jcp.10119 (acccessed 2024/03/25). Birben, E.; Sahiner, U. M.; Sackesen, C.; Erzurum, S.; Kalayci, O. Oxidative stress and antioxidant defense. World Allergy Organization Journal 2012, 5 (1), 9-19. DOI: 10.1097/WOX.0b013e3182439613. Martin, C.; Capilla-Lasheras, P.; Monaghan, P.; Burraco, P. The impact of chemical pollution across major life transitions: a meta-analysis on oxidative stress in amphibians. Proceedings of the Royal Society B: Biological Sciences 2024, 291 (2029), 20241536. DOI: 10.1098/rspb.2024.1536 (acccessed 2024/08/28).

(19) Angelier, F. Consequences of developmental exposure to pollution: Importance of stress-coping mechanisms. In Development Strategies and Biodiversity: Darwinian Fitness and Evolution in the Anthropocene, Springer, 2022; pp 283–316.

(20) Cuéllar-Valencia, O. M.; Murillo-García, O. E.; Rodriguez-Salazar, G. A.; Bolívar-García, W. Bioaccumulation of mercury in direct-developing frogs: The aftermath of illegal gold mining in a National Park. Herpetological Journal 2023, 33 (1), 6–13. Drummond, L. d. O.; Meire, R. O.; Braga, C.; Rezende, C. E. d.; Malm, O.; Cerqueira, R. Trophic position, altitudinal distribution, and water dependence as determining factors for mercury concentrations in tropical montane anurans. Science of The Total Environment 2022, 806, 151356. DOI: doi.org/10.1016/j.scitotenv.2021.151356. Loera, Y.; Gruppi, C.; Swing, K.; Campbell-Staton, S. C.; Milá, B.; Smith, T. B. Heavy metal contamination in birds from protected regions in the Amazon: heavy metals in birds from the Amazon. Environmental Toxicology and Chemistry 2024, 43 (12), 2601-2607. DOI: 10.1002/etc.5984 (acccessed 5/26/2026).

(21) Mittermeier, R. A.; Mittermeier, C. G.; Brooks, T. M.; Pilgrim, J. D.; Konstant, W. R.; da Fonseca, G. A. B.; Kormos, C. Wilderness and biodiversity conservation. Proceedings of the National Academy of Sciences 2003, 100 (18), 10309–10313. DOI: 10.1073/pnas.1732458100 (acccessed 2026/05/26).

(22) Guayasamin, J. M.; Ribas, C.; Carnaval, A.; Carrillo, J.; Hoorn, C.; Lohmann, L.; Riff, D.; Ulloa Ulloa, C.; Albert, J. Chapter 2: Evolution of Amazonian Biodiversity. In Amazon Assessment Report 2021, Nobre, C., Encalada, A., Anderson, E., Roca Alcazar, F., Bustamante, M., Mena, C., Peña-Claros, M., Poveda, G., Rodriguez, J., Saleska, S., et al. Eds.; United Nations Sustainable Development Solutions Network, 2021; pp 1–42.

(23) UNEP. Global mercury Assessment 2018; Geneva, Switzerland, 2019. Crespo-Lopez, M. E.; Augusto-Oliveira, M.; Lopes-Araújo, A.; Santos-Sacramento, L.; Yuki Takeda, P.; Macchi, B. d. M.; do Nascimento, J. L. M.; Maia, C. S. F.; Lima, R. R.; Arrifano, G. P. Mercury: What can we learn from the Amazon? Environment International 2021, 146, 106223. DOI: 10.1016/j.envint.2020.106223.

(24) Hödl, W. Reproductive diversity in Amazonian lowland frogs. Fortschritte der Zoologie 1990, 38, 41–60. Duellman, W. E. Reproductive Strategies of Frogs. Scientific American 1992, 267 (1), 80-87. (acccessed 2026/04/29/).JSTOR.

(25) Caldwell, J. P. Brazil nut fruit capsules as phytotelmata: interactions among anuran and insect larvae. Canadian Journal of Zoology-Revue Canadienne De Zoologie 1993, 71 (6), 1193–1201. Lehtinen, R. M.; Lannoo, M. J.; Wassersug, R. J. Phytotelm-breeding anurans: past, present, and future research. Miscellaneous Publications Museum of Zoology University of Michigan 2004, 193, 1-10. Summers, K.; McKeon, C. S. The evolutionary ecology of phytotelmata use in neotropical poison frogs. Miscellaneous Publications Museum of Zoology University of Michigan 2004, 193, 55-73.

(26) Haywood, L. K. Tadpoles as biological indicators of aquatic health: assessing the mining industries’ impact on water quality PhD Thesis, University of the Witwatersrand, Johannesburg, 2004. https://www.researchgate.net/profile/Lorren-Haywood/publication/326460927_Tadpoles_as_biological_indicators_of_aquatic_health_assessing_the_mining_industries’_impact_on_water_quality/links/5b4ed87f45851507a7a9ad2f/Tadpoles-as-biological-indicators-of-aquatic-health-assessing-the-mining-industries-impact-on-water-quality.pdf. Cuzziol Boccioni, Ana P.; Lajmanovich, Rafael C.; Repetti, María R.; Attademo, Andrés M.; Zalazar, Cristina S.; Manassero, A.; Russell-White, K.; Lancelle, María V.; Muchiutti, A.; Leon, Evelina J.; et al. Ecotoxicological impact of agro-industry on streams from a South American basin: amphibian tadpoles as indicators of environmental health. Water Environment Research 2025, 97 (6), e70117. DOI: 10.1002/wer.70117 (acccessed 2026/04/29).

(27) Luedtke, J. A.; Chanson, J.; Neam, K.; Hobin, L.; Maciel, A. O.; Catenazzi, A.; Borzée, A.; Hamidy, A.; Aowphol, A.; Jean, A.;, et al. Ongoing declines for the world’s amphibians in the face of emerging threats. Nature 2023, 622 (7982), 308–314. DOI: 10.1038/s41586-023-06578-4. Womack, M. C.; Steigerwald, E.; Blackburn, D. C.; Cannatella, D. C.; Catenazzi, A.; Che, J.; Koo, M. S.; McGuire, J. A.; Ron, S. R.; Spencer, C. L.; et al. State of the Amphibia 2020: A review of five years of amphibian research and existing resources. Ichthyology & Herpetology 2022, 110 (4), 638-661.

(28) Azambuja, G.; Kaefer, I.; Val, A. L.; Kochhann, D. A double threat to Amazonian amphibian larvae: a review of toxic contaminants and their interaction with global climate change. In The Future of Amazonian Aquatic Biota, de Souza, S. S., Braz-Mota, S., Val, A. L. Eds.; Springer Nature, 2024; pp 271–311. Silva, Y. B. d. S. e.; Ribeiro, B. R.; Thiesen Brum, F.; Soares-Filho, B.; Loyola, R.; Michalski, F. Combined exposure to hydroelectric expansion, climate change and forest loss jeopardies amphibians in the Brazilian Amazon. Diversity and Distributions 2018, 24 (8), 1072-1082. DOI: 10.1111/ddi.12745 (acccessed 2026/04/29).

(29) Buchman, M. NOAA screening quick reference tables, NOAA OR&R Report 08-1. Office of Response and Restoration Division, National Oceanic and Atmospheric Administration, Seattle 2008, 34.

(30) Wells, K. D. The Ecology and Behavior of Amphibians; University of Chicago Press, 2007. Duellman, W. E.; Trueb, L. Biology of Amphibians; McGraw Hill Publ. Co., 1994.

(31) Burke, J. N.; Bergeron, C. M.; Todd, B. D.; Hopkins, W. A. Effects of mercury on behavior and performance of northern two-lined salamanders (Eurycea bislineata). Environmental Pollution 2010, 158 (12), 3546–3551. DOI: 10.1016/j.envpol.2010.08.017.

(32) Bergeron, C. M.; Hopkins, W. A.; Todd, B. D.; Hepner, M. J.; Unrine, J. M. Interactive effects of maternal and dietary mercury exposure have latent and lethal consequences for amphibian larvae. Environmental Science & Technology 2011, 45 (8), 3781–3787. DOI: 10.1021/es104210a.

(33) Bergeron, C. M.; Hopkins, W. A.; Bodinof, C. M.; Budischak, S. A.; Wada, H.; Unrine, J. M. Counterbalancing effects of maternal mercury exposure during different stages of early ontogeny in American toads. Science of The Total Environment 2011, 409 (22), 4746–4752. DOI: 10.1016/j.scitotenv.2011.07.036. Unrine, J. M.; Jagoe, C. H. Dietary mercury exposure and bioaccumulation in southern leopard frog (Rana sphenocephala) larvae. Environmental Toxicology and Chemistry 2004, 23 (12), 2956-2963. DOI: doi.org/10.1897/03-695.1 (acccessed 2024/01/29).

(34) Born, M.; Gaucher, P. Distribution and life histories of amphibians and reptiles. In Nouragues: Dynamics and Plant-Animal Interactions in a Neotropical Rainforest., Bongers, F., Charles-Dominique, P., Forget, P.-M., Théry, M. Eds.; Vol. 80; Kluwer Academic Publisher, 2001; pp 167–184.

(35) Grant, T.; Frost, D. R.; Caldwell, J. P.; Gagliardo, R.; Haddad, C. F. B.; Kok, P. J. R.; Means, D. B.; Noonan, B. P.; Schargel, W. E.; Wheeler, W. C. Phylogenetic systematics of dart-poison frogs and their relatives (Amphibia: Athesphatanura: Dendrobatidae). Bulletin of the American Museum of Natural History 2006, 299, 6–262.

(36) Noonan, B. P.; Gaucher, P. Refugial isolation and secondary contact in the dyeing poison frog *Dendrobates tinctorius*. Molecular Ecology 2006, 15 (14), 4425–4435. DOI: 10.1111/j.1365-294X.2006.03074.x.

(37) Rojas, B. Strange parental decisions: fathers of the dyeing poison frog deposit their tadpoles in pools occupied by large cannibals. Behavioral Ecology and Sociobiology 2014, 68, 551–559. DOI: 10.1007/s00265-013-1670-y.

(38) Rojas, B.; Pašukonis, A. From habitat use to social behavior: natural history of a voiceless poison frog, Dendrobates tinctorius. PeerJ 2019, 7, e7648. DOI: 10.7717/peerj.7648.

(39) Ringler, E.; Pašukonis, A.; Hödl, W.; Ringler, M. Tadpole transport logistics in a Neotropical poison frog: indications for strategic planning and adaptive plasticity in anuran parental care. Frontiers in Zoology 2013, 10, 67.

(40) McKeon, C. S.; Summers, K. Predator driven reproductive behavior in a tropical frog. Evolutionary Ecology 2013, 27 (4), 725–737. DOI: 10.1007/s10682-013-9641-3.

(41) Bégué, L.; Tschirren, N.; Peignier, M.; Szabo, B.; Ringler, E. Behavioural consistency across metamorphosis in a neotropical poison frog. Evolutionary Ecology 2024, 38, 157–174. DOI: 10.1007/s10682-023-10274-0. Szabo, B.; Mangione, R.; Rath, M.; Pašukonis, A.; Reber, S. A.; Oh, J.; Ringler, M.; Ringler, E. Naive poison frog tadpoles use bi-modal cues to avoid insect predators but not heterospecific predatory tadpoles. Journal of Experimental Biology 2021, 224 (24), jeb243647.

(42) Jungfer, K.-H.; Faivovich, J.; Padial, J. M.; Castroviejo-Fisher, S.; Lyra, M. M.; V. M. Berneck, B.; Iglesias, P. P.; Kok, P. J. R.; MacCulloch, R. D.; Rodrigues, M. T.;, et al. Systematics of spiny-backed treefrogs (Hylidae: *Osteocephalus*): an Amazonian puzzle. Zoologica Scripta 2013, 42 (4), 351–380. DOI: 10.1111/zsc.12015 (acccessed 2026/06/12). Blotto, B. L.; Lyra, M. L.; Cardoso, M. C. S.; Trefaut Rodrigues, M.; R. Dias, I.; Marciano-Jr, E.; Dal Vechio, F.; Orrico, V. G. D.; Brandão, R. A.; Lopes de Assis, C.; et al. The phylogeny of the Casque-headed Treefrogs (Hylidae: Hylinae: Lophyohylini). Cladistics 2021, 37 (1), 36-72. DOI: doi.org/10.1111/cla.12409 (acccessed 2026/06/12).

(43) Jungfer, K.-H.; Weygoldt, P. Biparental care in tadpole-feeding Amazonian treefrog *Osteocephalus oophagus*. Amph.-Rept. 1999, 20, 235–249.

(44) Schiesari, L.; Gordo, M.; Hödl, W. Treeholes as calling, breeding, and developmental sites for the Amazonian canopy frog, Phrynohyas resinifictrix (Hylidae). Copeia 2003, 2003 (2), 263–272.

(45) Mignet, F. Biology and captive breeding of the Amazonian milk frog, *Trachycephalus resinifictrix* (Goeldi 1907) *Reptiles & Amphibians*, 2015, 22 (2), 68–75. DOI: doi.org/10.17161/randa.v22i2.14047.

(46) Caldwell, J. P. A new species of toad in the genus *Bufo* from Pará, Brazil, with an unusual breeding site. Pap. Avul. Zool. Mus. Zool. Univ. São Paulo 1991, 37, 389–400.

(47) de Figueiredo, V. A.; dos Santos, E. M.; Pedroso-Santos, F.; Sanches, P. R.; Costa-Campos, C. E. Diet and clutch size of Rhinella castaneotica (Anura: Bufonidae) from a forest area in Serra do Navio, Amapá, Brazil. North-Western Journal of Zoology 2022, 18, 35–39.

(48) Protázio, A. d. S.; Gama, V.; Silva, S. V.; dos Santos, C. G. C.; de Oliveira, J. K. G. Diet of tadpoles for five anuran species of northeast Brazil. Journal of Limnology 2020, 79, 180–186. DOI: doi.org/10.4081/jlimnol.2020.1912. Bionda, C.; Gari, N.; Luque, E.; Salas, N.; Lajmanovich, R.; Martino, A. Ecología trófica en larvas de Rhinella arenarum (Anura: Bufonidae) en agroecosistemas y sus posibles implicaciones para la conservación. Revista de Biología Tropical 2012, 60, 771-779.

(49) Abràmoff, M. D.; Magalhães, P. J.; Ram, S. J. Image processing with ImageJ. Biophotonics International 2004, 11 (7), 36–42.

(50) Gosner, K. L. A simplified table for staging anuran embryos and larvae with notes on identification. Herpetologica 1960, 16, 183–194.

(51) Zuur, A. F.; Ieno, E. N.; Walker, N. J.; Saveliev, A. A.; Smith, G. M. Mixed effects models and extensions in ecology with R; Springer, 2009.

(52) emmeans: Estimated Marginal Means, aka Least-Squares Means 2026. (accessed.

(53) Pike, N. Using false discovery rates for multiple comparisons in ecology and evolution. Methods in Ecology and Evolution 2011, 2 (3), 278–282. DOI: 10.1111/j.2041-210X.2010.00061.x (acccessed 2026/06/12).

(54) Bates, D.; Maechler, M.; Bolker, B.; Walker, S. Fitting Linear Mixed-Effects Models Using lme4. Journal of Statistical Software 2015, 67, 1–48. DOI: doi:10.18637/jss.v067.i01.

(55) DHARMa: Residual Diagnostics for Hierarchical (Multi-Level / Mixed) Regression Models; 2024. (accessed.

(56) RCoreTeam. R: A language and environment for statistical computing. R Foundation for Statistical Computing, Vienna, Austria. 2013.

(57) Wang, Q.; Kim, D.; Dionysiou, D. D.; Sorial, G. A.; Timberlake, D. Sources and remediation for mercury contamination in aquatic systems—a literature review. Environmental Pollution 2004, 131 (2), 323–336. DOI: 10.1016/j.envpol.2004.01.010. Sagisaka Méndez, S.; Mitchell, C. P. J. Effects of temperature on mercury methylation and demethylation in Boreal wetland soils. Journal of Geophysical Research: Biogeosciences 2026, 131 (5), e2025JG009302. DOI: doi.org/10.1029/2025JG009302 (acccessed 2026/06/11).

(58) Regnell, O. The effect of pH and dissolved oxygen levels on methylation and partitioning of mercury in freshwater model systems. Environmental Pollution 1994, 84 (1), 7–13. DOI: 10.1016/0269-7491(94)90064-7.

(59) Cleophas, F.; Mahali, M.; Musta, B.; Zahari, N. Z.; Lee, L. E.; Bidin, K. Canopy precipitation interception in a lowland tropical forest in relation to stand structure. IOP Conference Series: Earth and Environmental Science 2022, 1053 (1), 012005. DOI: 10.1088/1755-1315/1053/1/012005.

(60) Bank, M. S.; Crocker, J.; Connery, B.; Amirbahman, A. Mercury bioaccumulation in green frog (*Rana clamitans*) and bullfrog (*Rana catesbeiana*) tadpoles from Acadia National Park, Maine, USA. Environmental Toxicology and Chemistry 2007, 26 (1), 118–125.

(61) Loftin, C., S.; Calhoun, A., J. K.; Nelson, S., J.; Elskus, A., A.; Simon, K. Mercury bioaccumulation in wood frogs developing in seasonal pools. Northeastern Naturalist 2012, 19 (4), 579–600. DOI: 10.1656/045.019.0404.

(62) Regnell, O.; Watras, C. J. Microbial mercury methylation in aquatic environments: a critical review of published field and laboratory studies. Environmental Science & Technology 2019, 53 (1), 4–19. DOI: 10.1021/acs.est.8b02709.

(63) Unrine, J. M.; Hopkins, W. A.; Romanek, C. S.; Jackson, B. P. Bioaccumulation of trace elements in omnivorous amphibian larvae: Implications for amphibian health and contaminant transport. Environmental Pollution 2007, 149 (2), 182–192. DOI: 10.1016/j.envpol.2007.01.039. Unrine, J. M.; Jagoe, C. H.; Brinton, A. C.; Brant, H. A.; Garvin, N. T. Dietary mercury exposure and bioaccumulation in amphibian larvae inhabiting Carolina bay wetlands. Environmental Pollution 2005, 135 (2), 245-253. DOI: doi.org/10.1016/j.envpol.2004.11.003.

(64) Snodgrass, J. W.; Hopkins, W. A.; Jackson, B. P.; Baionno, J. A.; Broughton, J. Influence of larval period on responses of overwintering green frog (*Rana clamitans*) larvae exposed to contaminated sediments. Environmental Toxicology and Chemistry 2005, 24 (6), 1508–1514. DOI: 10.1897/04-339R1.1 (acccessed 2026/05/30).

(65) St. Louis, V. L.; Rudd, J. W. M.; Kelly, C. A.; Hall, B. D.; Rolfhus, K. R.; Scott, K. J.; Lindberg, S. E.; Dong, W. Importance of the forest canopy to fluxes of methyl mercury and total mercury to Boreal ecosystems. Environmental Science & Technology 2001, 35 (15), 3089–3098. DOI: 10.1021/es001924p.

(66) Rojas, B. Mind the gap: treefalls as drivers of parental tradeoffs. Ecology and Evolution 2015, 5 4028–4036. DOI: doi: 10.1002/ece3.1648. Fouilloux, C. A.; Fromhage, L.; Valkonen, J. K.; Rojas, B. Size-dependent aggression towards kin in a cannibalistic species. Behavioral Ecology 2022, 33, 582–591. DOI: 10.1093/beheco/arac020 (acccessed 3/27/2022).

(67) Lemaire, J.; Mangione, R.; Caut, S.; Bustamante, P. Mercury biomagnification in the food web of Agami Pond, Kaw-Roura Nature Reserve, French Guiana. Heliyon 2024, 10 (7). DOI: 10.1016/j.heliyon.2024.e28859 (acccessed 2025/03/24).

(68) Davidson, M. A.; Croteau, M. C.; Millar, C. S.; Trudeau, V. L.; Lean, D. R. S. Fate and developmental effects of dietary uptake of methylmercury in *Silurana tropicalis* tadpoles. Journal of Toxicology and Environmental Health, Part A 2011, 74 (6), 364–379. DOI: 10.1080/15287394.2011.534427.

(69) Bergeron, C. M.; Bodinof, C. M.; Unrine, J. M.; Hopkins, W. A. Bioaccumulation and maternal transfer of mercury and selenium in amphibians. Environmental Toxicology and Chemistry 2010, 29 (4), 989–997, 10.1002/etc.125. DOI: doi.org/10.1002/etc.125 (acccessed 2023/02/13). Lemaire, J.; Marquis, O.; Bustamante, P.; Mangione, R.; Brischoux, F. I got it from my mother: Inter-nest variation of mercury concentration in neonate Smooth-fronted Caiman (Paleosuchus trigonatus) suggests maternal transfer and possible phenotypical effects. Environmental Research 2021, 194, 110494. DOI: doi.org/10.1016/j.envres.2020.110494.

(70) Ackerman, J. T.; Herzog, M. P.; Evers, D. C.; Cristol, D. A.; Kenow, K. P.; Heinz, G. H.; Lavoie, R. A.; Brasso, R. L.; Mallory, M. L.; Provencher, J. F.;, et al. Synthesis of maternal transfer of mercury in birds: implications for altered toxicity risk. Environmental Science & Technology 2020, 54 (5), 2878–2891. DOI: 10.1021/acs.est.9b06119.

(71) Saxton, H. J.; Goodman, J. R.; Collins, J. N.; Black, F. J. Maternal transfer of inorganic mercury and methylmercury in aquatic and terrestrial arthropods. Environmental Toxicology and Chemistry 2013, 32 (11), 2630–2636. DOI: 10.1002/etc.2350 (acccessed 5/29/2026).

(72) Ackerman, J. T.; Eagles-Smith, C. A.; Herzog, M. P.; Hartman, C. A. Maternal transfer of contaminants in birds: Mercury and selenium concentrations in parents and their eggs. Environmental Pollution 2016, 210, 145–154. DOI: 10.1016/j.envpol.2015.12.016.

(73) Yang, L.; Zhang, Y.; Wang, F.; Luo, Z.; Guo, S.; Strähle, U. Toxicity of mercury: Molecular evidence. Chemosphere 2020, 245, 125586. DOI: 10.1016/j.chemosphere.2019.125586. Wu, Y.-S.; Osman, A. I.; Hosny, M.; Elgarahy, A. M.; Eltaweil, A. S.; Rooney, D. W.; Chen, Z.; Rahim, N. S.; Sekar, M.; Gopinath, S. C. B.; et al. The toxicity of mercury and its chemical compounds: molecular mechanisms and environmental and human health implications: a comprehensive review. ACS Omega 2024, 9 (5), 5100-5126. DOI: 10.1021/acsomega.3c07047.

(74) Tettamanti, G.; Casartelli, M. Cell death during complete metamorphosis. Philosophical Transactions of the Royal Society B: Biological Sciences 2019, 374 (1783), 20190065. DOI: 10.1098/rstb.2019.0065 (acccessed 10/9/2026).

(75) Su, Y.; Damjanovski, S.; Shi, Y.; Shi, Y. Molecular and cellular basis of tissue remodeling during amphibian metamorphosis. Histology and Histopathology 1999, 14 (1), 175–183. DOI: 10.14670/hh-14.175

(76) Marasco, V.; Stier, A.; Boner, W.; Griffiths, K.; Heidinger, B.; Monaghan, P. Environmental conditions can modulate the links among oxidative stress, age, and longevity. Mechanisms of Ageing and Development 2017, 164, 100–107. DOI: 10.1016/j.mad.2017.04.012. Dai, L.; Watanabe, M.; Qureshi, A. R.; Mukai, H.; Machowska, A.; Heimbürger, O.; Barany, P.; Lindholm, B.; Stenvinkel, P. Serum 8-hydroxydeoxyguanosine, a marker of oxidative DNA damage, is associated with mortality independent of inflammation in chronic kidney disease. European Journal of Internal Medicine 2019, 68, 60-65. DOI: doi.org/10.1016/j.ejim.2019.07.035.

